# Neuro-immunobiology and treatment assessment in a mouse model of anti-NMDAR encephalitis

**DOI:** 10.1101/2024.07.19.604240

**Authors:** Estibaliz Maudes, Jesús Planagumà, Laura Marmolejo, Marija Radosevic, Ana Beatriz Serafim, Esther Aguilar, Carlos Sindreu, Jon Landa, Anna García-Serra, Francesco Mannara, Marina Cunquero, Anna Smith, Chiara Milano, Paula Peixoto-Moledo, Mar Guasp, Raquel Ruiz-Garcia, Sarah M. Gray, Marianna Spatola, Pablo Loza-Alvarez, Lidia Sabater, Carlos Matute, Josep Dalmau

**Author notes:** **Correspondence to:** Josep Dalmau, Neuroimmunology Program, Fundació Clinic per la Recerca Biomèdica – Institut d’Investigacios Biomèdiques August Pi i Sunyer (FCRB-IDIBAPS), University of Barcelona, Casanova, 143; CELLEX3A, Barcelona 08036, Spain. These authors contributed equally to this work (Jesús Planagumà, Laura Marmolejo).

## Abstract

Anti-N-methyl-*D*-aspartate receptor (NMDAR) encephalitis is a disorder mediated by autoantibodies against the GluN1 subunit of NMDAR. It occurs with severe neuropsychiatric symptoms that often improve with immunotherapy. Clinical studies and animal models based on patients’ antibody transfer or NMDAR immunization suggest that the autoantibodies play a major pathogenic role. Yet, there is an important need of models offering an all-inclusive neuro-immunobiology of the disease together with a clinical course long enough to facilitate the assessment of potential new treatments. Toward this end, eight-week-old female mice (C57BL/6J) were immunized (days 1 and 28) with GluN1_356-385_ peptide or saline with AddaVax adjuvant and pertussis toxin. After symptom development (∼day 35), subsets of mice were treated with an anti-CD20 (day 35), a positive allosteric modulator (PAM) of NMDAR (NMDAR-PAM, SGE-301) from days 45 to 71, or both. GluN1-antibody synthesis, epitope spreading, effects of antibodies on density and function of NMDAR, brain immunological infiltrates, microglial activation and NMDAR phagocytosis, and antibody synthesis in cultured inguinal and deep cervical lymph nodes (DCLN) were assessed with techniques including immunohistochemistry, calcium imaging, confocal and super-resolution microscopy, electrophysiology, or flow cytometry. Changes of memory and behaviour were assessed with a panel of behavioural tests, and clinical/subclinical seizures with brain-implanted electrodes. Immunized mice, but not controls, developed serum and CSF NMDAR-antibodies (IgG1 predominant) against the immunizing peptide and other GluN1 regions (epitope spreading) resulting in a decrease of synaptic and extrasynaptic NMDAR clusters and reduction of hippocampal plasticity. These findings were associated with brain inflammatory infiltrates, mainly B- and plasma cells, microglial activation, colocalization of NMDAR-IgG complexes with microglia, and presence of these complexes within microglial endosomes. Cultures of DCLC showed GluN1-antibody production. These findings were associated with psychotic-like behaviour (predominant at disease onset), memory deficit, depressive-like behaviour, abnormal movements (15% of mice), and lower threshold for developing pentylenetetrazole-induced seizures (hypoactivity, myoclonic jerks, continuous tonic-clonic) which correlated with regional cFOS expression. Most symptoms and neurobiological alterations were reversed by the anti-CD20 and PAM, alone or combined. Initial repopulation of B cells, by the end of the study, was associated with re-emergence of clinical-neurobiological alterations, which were abrogated by PAM. Overall, this model offers an all-inclusive neuro-immunobiology of the disease, allowing testing novel treatments, supporting the potential therapeutic role of NMDAR-PAM, and suggesting an immunological paradigm of systemic antigen presentation and brain NMDAR epitope spreading, which along the DCLN might contribute to fine-tune the polyclonal immune response.

## Introduction

Anti-N-methyl-*D*-aspartate receptor (NMDAR) encephalitis is an autoimmune brain disease of rapid presentation and prolonged clinical course characterized by severe psychiatric and neurological symptoms in association with IgG antibodies against the GluN1 subunit of NMDARs.^1^ Most patients are young adults and children, predominantly female (4:1), who in a matter of days or weeks develop behavioural change, psychosis, memory impairment, and variable presence of abnormal movements, seizures, decreased level of consciousness or dysautonomia.^2^ Known triggers of the disease are tumors, mainly teratomas, and less frequently herpes simplex encephalitis, but in about 60% of patients the trigger is unknown.^3, 4^ Treatments aimed to remove the antibodies and antibody-producing cells (e.g., anti-CD20 such as rituximab) often result in improvement, but for most patients the recovery is slow, remaining with memory and cognitive deficits for several months.^5^ Because the pathophysiology of the post-acute stage is unknown, the treatment approach to shorten the period of recovery and improve clinical outcome is currently one of the most important challenges of the disease.^6^

In cultures of rodent hippocampal neurons, patients’ antibodies disrupt the surface dynamics of NMDARs and internalize them, causing a reduction of their surface content and NMDAR-mediated currents.^7, 8^ Similar effects are obtained with passive transfer of patients’ antibodies to the cerebroventricular system of mice, causing transient psychotic-like behaviour, memory impairment, and seizures or reduced seizure threshold.^9–11^ These studies confirm the pathogenicity of patients’ autoantibodies but do not represent a *bona fide* model of anti-NMDAR encephalitis, and are unable to provide further insights into the immunopathology and clinical course of the disease.

Additional approaches to model the disease are based on immunization of rodents with native NMDAR or peptides of the GluN1 subunit, resulting in phenotypes that range from fulminant, sometimes lethal, encephalitis (with severe seizures and stereotyped movements) to milder phenotypes that depending on the model, include memory impairment, behavioural change (depressive, anxiety-like), or reduction of seizure threshold.^12–17^ These models occur with the development of NMDAR antibodies and support the concept that NMDAR autoimmunity is sufficient to cause multifaceted symptoms. However, there is an unmet need of models that can offer an all-inclusive assessment of the neurobiology and immunobiology of the disease and a clinical course long enough to facilitate the evaluation of potential therapies on all these paradigms.

Even though animal models are imperfect reproductions of human diseases, they offer insights into the pathogenic mechanisms and potential new treatments, which for anti-NMDAR encephalitis is of paramount importance. Towards this end, we developed a mouse model of anti-NMDAR encephalitis that allows behavioural, neurobiological and immunobiological assessments during an extended course. Additionally, we tested the model with several treatment approaches including an anti-CD20, which represents a frequent treatment in patients; a positive allosteric modulator (PAM) of NMDAR (SGE-301) that has potential for clinical use,^18, 19^ and both combined, and determined how these treatments modified the clinical and biological paradigms of the disease.

## Materials and methods

### Animals

Female C57BL6/J mice (Charles River) were housed in cages of four in our animal facility (Unitat d’Experimentació Animal de Medicina, Centres Científics i Tecnològics, Universitat de Barcelona) at a controlled temperature (21 ± 1°C) and humidity (55 ± 10%) with illumination at 12-h cycles, and food and water *ad libitum*. Experiments were performed during the light phase, and animals were habituated to the room for 30 min before each experiment. All procedures were done according to standard ethical guidelines (European Communities Directive 2010/63/EU), approved by the local ethical committee (CEEA-316/22), and reported in accordance with the ARRIVE guidelines (Supplementary Material).

### Immunization and treatments

On days 1 and 28, eight-week-old mice were subcutaneously injected with 200 µg of the GluN1_356–385_ peptide or saline and AddaVax (InvivoGen, San Diego, USA) adjuvant. All animals received 100 ng of Bordetella pertussis toxin intraperitoneally at the time of immunization and 48 h later. To assess different treatments, a subset of mice received 250 µg of anti-CD20 intravenously on day 35, another subset received from day 47 until 71 (end of the study) daily intraperitoneal injections of a NMDAR-PAM (SGE-301), and a third subset received both, anti-CD20 and PAM (Fig. 1A). Each subset had the corresponding controls including intravenous injection of saline, intraperitoneal injection of vehicle, or both. A total of 275 animals were used in the study. Mice were randomly allocated to NMDAR or control groups. The preparation and concentration of SGE-301 (10 mg/kg) were similar as those previously reported.^18^

**Figure 1:**
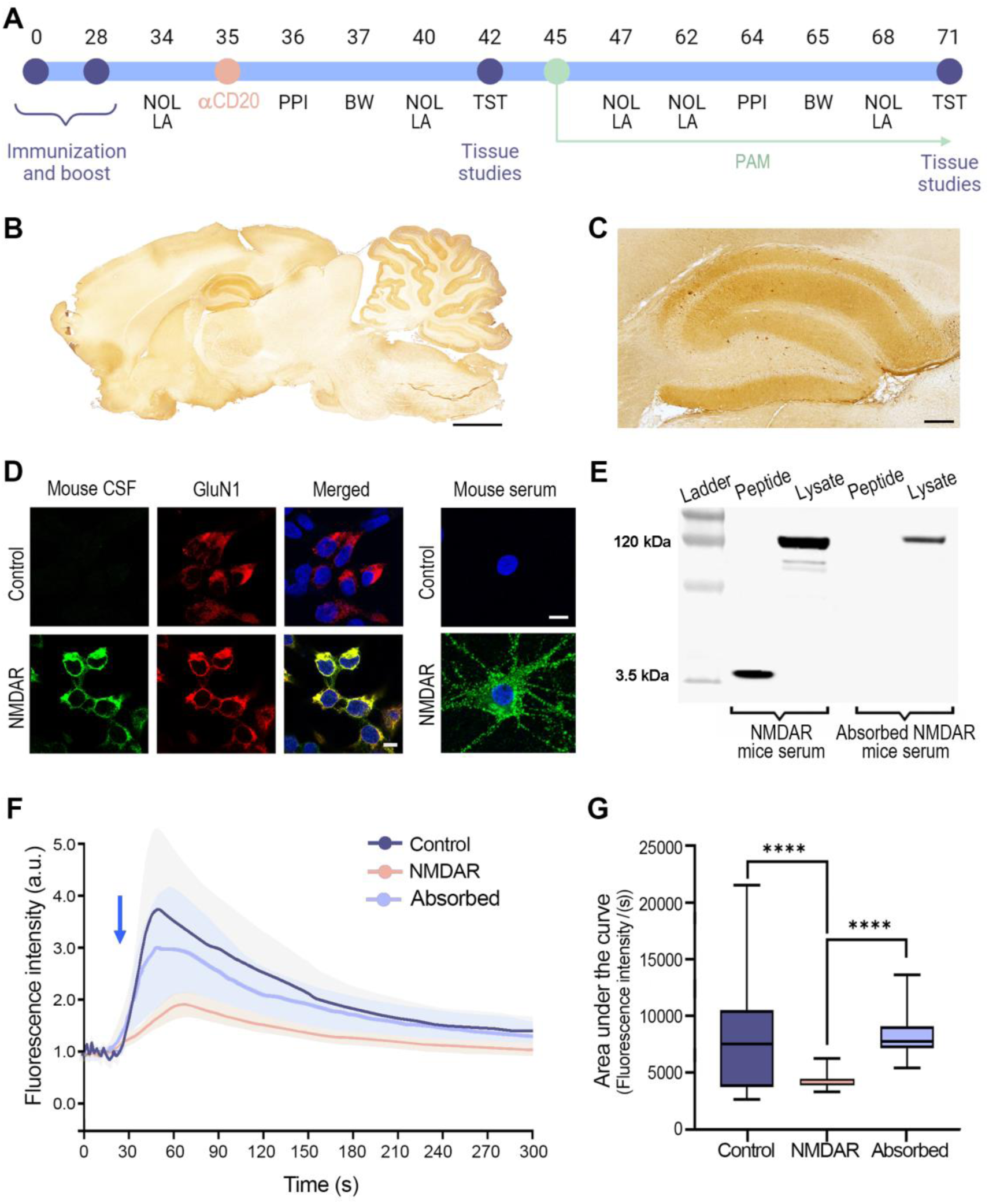
Immunization with GluN1_356-385_ results in pathogenic GluN1 antibody production. **(A)** Schematic representation of the immunization, treatment, behavioural tasks and tissue study timeline. NOL: Novel Object Location; PPI: Prepulse Inhibition; BM: Black and White test; TST: Tail Suspension Test. On days 42 and 71 subsets of mice were sacrificed and their blood, spleen, lymph nodes, brain, and CSF were harvested for tissue and cellular studies. **(B)** Sagittal section of rat brain immunostained with IgG purified from a pool of 5 NMDAR mice showing a pattern of neuropil staining similar to that described for patients’ NMDAR IgG. Scale bar = 2 mm. **(C)** Magnification showing the immunostaining of the hippocampus. Scale bar = 250 µm. **(D)** Left side: cell-based assay with HEK293 cells expressing GluN1/GluN2b showing reactivity with CSF from an NMDAR mouse (green), which colocalizes (yellow) with the commercial GluN1 antibody (red); CSF from a control mouse shows no reactivity. Right side: live neuron immunofluorescence showing intense cell-surface immunolabelling (green) with serum of an NMDAR mouse, but not with control serum. **(E)** Immunoblot of GluN1_356-385_ peptide and neuronal lysate probed with non-absorbed and peptide-absorbed NMDAR mice serum. Compared with non-absorbed NMDAR mice serum, the peptide-absorbed mice serum is no longer reactive with the peptide, but retains reactivity with the neuronal lysate. **(F)** NMDAR mice IgG reduce NMDA-induced calcium influx in rat neurons expressing GCaMP5G. The plot shows one of three independent experiments representing the fluorescence intensity upon NMDA stimulation (blue arrow) for neuronal cultures treated with NMDAR mice IgG (pink), control mice IgG (dark blue), and NMDAR mice IgG pre-absorbed with NMDAR (light blue). Data are represented as mean ± SEM. **(G)** Analysis of the area under the curve of calcium fluorescence over-time for the three experimental groups (controls n = 56 cells; NMDAR mice n = 103 cells; NMDAR mice absorbed n = 43 cells). Box plots show the median, 25^th^ and 75^th^ percentile. Whiskers indicate the minimum and maximum values. Assessment of significance was performed by one-way analysis of variance (ANOVA, p < 0.0001). A value of p < 0.05 was considered statistically significant.

### GluN1 antibody detection and characterization

Serum was obtained from blood collected from the submandibular vein, and CSF from the cisterna magna of deeply anesthetized mice prior to sacrifice on day 42 or 71. The presence of GluN1 antibodies was determined with rat brain immunohistochemistry, cell-based assay (CBA) of GluN1/GluN2b, and live immunolabeling of cultures of dissociated rat hippocampal neurons, as reported.^2, 8^ The titre of NMDAR antibodies was calculated by CBA with serial dilutions of samples until the reactivity was no longer visible (units reported as dilution factor). Mice NMDAR antibody class and subclass were determined with the indicated CBA of GluN1/GluN2b and antibodies against each subclass of mouse IgG (IgG1, 2, 3) (Supplementary material).

The occurrence of antibodies targeting epitope regions other than that of the immunizing GluN1_356–385_ peptide (epitope spreading) was determined with CBA of a truncated GluN1 construct without the peptide sequence,^20^ and with western blot of the immunizing peptide and neuronal lysates probed with mice serum non-absorbed and pre-absorbed with the immunizing peptide (Supplementary Material).

A preliminary assessment of functional effects of IgG from GluN1 immunized mice (NMDAR mice) was conducted using quantitative immunocytochemical analysis of clusters of NMDARs in live rat hippocampal neurons exposed for 24 h to IgG of NMDAR mice or controls, and calcium imaging of similar cultures of neurons under the same experimental conditions (Supplementary Material). The specificity of the NMDAR-IgG effect in calcium imaging experiments was determined with IgG pre-absorbed with GluN1-expressing HEK293 cells. The techniques of immunocytochemical quantitation of NMDAR clusters, IgG isolation, and immunoabsorption have been previously reported,^8, 21, 22^ and the calcium imaging experiment is described in Supplementary Material.

Determination of IgG presence in the brain, immunoprecipitation of IgG bound to NMDAR, and assessment of complement deposition was assessed as previously reported and described in Supplementary Material.^23^

### Brain, spleen, and lymph node dissection

On days 42 and 71, subsets of mice were deeply anaesthetized, and their spleen was harvested. Mice were then euthanized by cardiac perfusion with saline, and the brain was removed. For immunohistochemical and confocal microscopy studies, the right hemisphere was fixed with 4% paraformaldehyde for 1h, cryopreserved in 40% glucose for 48h, embedded in optimal cutting temperature compound, and snapped frozen in isopentane chilled with liquid nitrogen. The left hemisphere was fresh frozen for immunoprecipitation studies, acutely sectioned for electrophysiology, or processed to obtain immune cells for flow cytometry.

Inguinal and deep cervical lymph nodes of immunized mice and controls were obtained at day 42. Lymph node cells were dissociated and kept in culture with X-vivo 15 (02-053Q, Lonza, Basel, Switzerland) supplemented with 10% fetal bovine serum for 24 h, and with the presence of 0,1 µg/µl of GluN1_356–385_ peptide for 3 days. The media was then assessed (diluted 1:2) for the presence of GluN1 antibodies with a cell-based assay (CBA) expressing GluN1/GluN2b.

### Analysis of brain and spleen immune cells

#### Flow cytometry

Brain tissue was homogenized with gentleMACS Dissociator (130-096-427, Miltenyi Biotec) while immersed in 2ml of HBSS buffer (w/o Ca^2+^ and Mg^2+,^ 14175-053, Thermo Fisher Scientific) containing 100 U/mL collagenase IV (C5138, Sigma) and 50 U/mL DNase I (D5025-150 KU, Sigma). The tissue was then filtered on a cell strainer (70 µm) and the cells were separated from myelin and debris with a 30% Percoll gradient (17-0891-01, GE Healthcare) in HBSS without Ca^2+^ and Mg^2+^, centrifuged at 950 g during 25 min without brakes. Cells were collected from the bottom of the tube after centrifugation and washed with HBSS buffer. The isolation of immune cells from the spleen is provided in Supplementary Material.

Brain immune cell infiltrates or splenocytes were incubated with the following fluorophore-conjugated antibodies during 20 min at 4 °C: CD11b (clone M1/70, eFluor 450, 48011282), CD45 (clone 30-F11, APC, 17045182), CD3 (clone 17A2, Alexa Fluor 488, 53003282), CD4 (clone RM4-5, PE-Cyanine5.5, 35004282), CD8 (clone 53-6.7, Brilliant Ultra Violet 737, 367008182), CD19 (clone 1D3, PE-Cyanine5, 15019382), IgD (clone 11-26, Super Bright 600, 63599382), CD27 (clone LG.7F9, Super Bright 702, 67027182), CD138 (clone 300506, PE, MA523527). All the antibodies were purchased from Invitrogen (Waltham, MA, USA). Data was acquired in a Cytek Aurora cytometer and analysed using SpectroFlo software (Cytek Biosciences, Fremont, CA, USA).

#### Brain immunohistochemistry

Additional studies of brain T cell infiltrates were performed by immunostaining using antibodies specific for CD4 and CD8 T cells as described in Supplementary Material.

#### ELISpot

Determination of GluN1_356–385_ specific T cells in splenocytes was performed with ELISpot, described in Supplementary Material.

### NMDAR cluster density and microglia studies

To determine the cluster density of cell-surface NMDAR and postsynaptic density protein 95 (PDS95), 5 µm-thick brain sections of NMDAR mice and controls were incubated with a human CSF enriched with GluN1 antibodies (used as a primary antibody) for 1h at room temperature (RT), followed by the secondary Alexa Fluor 488 goat anti-human IgG (1:1000, A-11013, Thermo Fisher) for 1h at RT, as reported.^9^ Tissue sections were then permeabilized with 0.3% Triton X-100 for 10 min at RT and incubated with rabbit polyclonal anti-PSD95 (1:250, ab18258, Abcam, Cambridge, UK) overnight at 4°C, followed by the corresponding secondary Alexa Fluor 594 goat anti-rabbit IgG (1:1000, A-11012, Thermo Fisher) for 1h at RT.

Microglia was assessed in brain sections using a monoclonal rat antibody against CD68 (1:200, MCA1957GA, Bio-Rad, Hercules, CA, USA) to label macrophage/microglia and a polyclonal rabbit antibody against Iba-1 (1:1000, 019-19741, Wako Chemicals, Neuss, Germany) to label activated microglia. To assess whether microglia co-localized with IgG bound to NMDAR, we used a triple staining (anti-CD68, anti-mouse IgG, anti-NMDAR) each as indicated above, for 2h at RT, followed by the corresponding secondary antibodies for 1h at RT: goat anti-rat Alexa Fluor 594 (1/500, A-1100), goat anti-mouse Alexa Fluor 488 (1:500, A-11001), and goat anti-human Alexa Fluor 647 (1:500, A-21445) all from Thermo Fisher (Waltham, USA).

Slides were then mounted in ProLong Gold antifade (P36935, Thermo Fisher) and scanned under a Zeiss LSM710 confocal microscope (Carl Zeiss, Jena, Germany) with the EC-Plan NEOFLUAR CS 100x/1.3 NA oil objective. Cluster analysis was performed as reported.^9^ In brief, standardized z-stacks including 50 optical images were acquired from the CA1, CA3, dentate gyrus (DG) and cortex. Images were then deconvolved using Huygens Essential 23.10 software (Scientific Volume Imaging, Hilversum, NL), and a spot detection algorithm from Imaris 8.1 software (Oxford instruments, Belfast, UK) was used. Density of clusters was expressed as spots/µm^3^. Three-dimensional colocalization of clusters was done using a spot colocalization algorithm (Imaris 8.1, Oxford instruments). Synaptic localization was defined as colocalization of NMDAR with PSD95. Microglia activation was defined as colocalization of Iba-1 with CD68. Phagocytosis of the complex IgG-NMDAR was defined by the triple colocalization of CD68, mouse IgG, and NMDAR, and confirmed with Stimulated Emission Depletion (STED) microscopy (Supplementary Material).

### Hippocampal long-term potentiation, and paired-pulse facilitation

Acute sections of the hippocampus on day 42 and 71 were used to assess long-term plasticity by the classical paradigm of stimulation at the Schaffer collateral pathway and recording the field potentials at CA3-CA1 synapses, as previously described.^18, 19^ A detailed description can be found in Supplementary Material.

### Behavioural testing, seizure susceptibility, and abnormal movements

A panel of standardized behavioural and memory tests was applied by investigators blinded to the experimental conditions (Fig. 1A). They included: memory (Novel Object Location [NOL]), psychotic-like behaviour (Pre-Pulse Inhibition [PPI]), anxiety (Black and White [BW]), depressive-like behaviour (Tail Suspension Test [TST]), and locomotor activity (LA). These tests are described in Supplementary Material and have been reported.^9, 10^ Mice were subjected to PPI, BW and TST just once prior to being sacrificed on day 42 or 71. Mice sacrificed on day 71 underwent NOL test from the beginning of the battery. To determine seizure susceptibility, the GABA_a_R antagonist pentylenetetrazol (PTZ, Sigma) was given intraperitoneally (40 mg/kg) to a subgroup of NMDAR and control mice on days 38-43. Seizure development and scaling was confirmed by recordings via intracerebral electrodes synchronized to a video camera, and by cFos immunostaining (Supplementary Material). The presence of abnormal movements was visually assessed, without quantification, during recordings or the indicated behavioural tests.

### Statistical analysis

Comparisons of total and synaptic NMDAR clusters, IgG deposits, field excitatory postsynaptic potentials (fEPSP) slope change, brain infiltrates, splenic B cells, T cell activation, microglia activation, and behavioural tests across of all treatment groups were conducted using a mixed-effects model. This model included immunization, treatment, and time as fixed effects, with the subject included as a random effect to account for inter-subject variability. The area under the curve of calcium curves was also analyzed with a mixed-effects model, where IgG type was a fixed effect and hippocampal culture a random effect to account for culture replicate viability. These analyses were performed using the lme4 package in R, with p-values adjusted for multiple comparisons using Bonferroni correction. Comparison of hippocampal NMDAR and PSD95 clusters, and microglia phagocytosis were performed using a nested t-test. PPI test, BW test and TST for untreated control and NMDAR mice were compared using a two-way analysis of variance with Bonferroni correction for multiple comparisons. Seizure susceptibility was evaluated using a chi-squared test. All experiments were assessed for outliers with the ROUT method applying Q = 1%. In all analyses, we used a 2-sided type I error of 5%. All tests and graphs were performed using GraphPad Prism (version 8; GraphPad Inc., San Diego, CA) and R studio (v4.0.0).

### Data availability

The data that support the findings of this study are available from the corresponding author, upon request.

## Results

### NMDAR mice produce GluN1 antibodies with effects similar to those of anti-NMDAR encephalitis

NMDAR mice, but not controls, developed serum and CSF NMDAR antibodies that were demonstrated with rat brain immunostaining (Fig. 1B-C), CBA with HEK293 cells expressing GluN1/GluN2B subunits of NMDAR, and live immunolabelling of cultures of rat hippocampal neurons (Fig. 1D). The predominant GluN1 immunoglobulin class and subclass was examined in a representative group of 10 mice, showing IgG1 in all, and IgG2 in 4 (40%) (data not shown). Immunoabsorption of the antibodies with GluN1_356–385_ peptide abrogated the reactivity with the peptide but not with GluN1 regions outside the peptide sequence, suggesting epitope spreading (Fig. 1E); this was confirmed with CBA of a GluN1 construct that did not contain the peptide sequence (Supplementary Fig. 1). In cultured rat hippocampal neurons, IgG from NMDAR mice, but not from controls, caused a significant reduction of cell-surface NMDAR clusters (Supplementary Fig. 2). Moreover, neurons pre-treated with IgG from NMDAR mice showed a significant reduction of NMDA-induced calcium influx compared with neurons pre-treated with IgG from controls. This effect was abrogated if the IgG from NMDAR mice had been pre-absorbed with NMDARs (Fig. 1F-G, Supplementary Video 1). Overall, these findings show that NMDAR mice developed polyclonal antibodies highly similar in all paradigms tested to those reported in the human disease.^8, 9, 24^ Moreover, the presence of GluN1 antibodies was confirmed in the media of cultured cells from deep cervical lymph nodes, but not inguinal lymph nodes, of NMDAR mice (Supplementary Fig. 3).

### Preliminary assessment of anti-CD20 and PAM treatments used in the model

In previous studies with passive transfer of patients’ NMDAR antibodies to mice, and in cultured neurons, we previously established the effects of the NMDAR-PAM (SGE-301).^18, 19, 25^ Since we had not previously tested the anti-CD20 used in the current model, we preliminary confirmed that it was able to deplete systemic B cells in mice. Flow cytometry on splenocytes from all subsets of NMDAR and control mice showed that administration of anti-CD20 at day 35 caused a significant reduction of B cells measured at day 42, which was no longer detected by day 71, indicating B cell repopulation (Supplementary Fig. 4A-B).

Additional studies with IFN-γ ELISpot on splenocytes from NMDAR mice showed activation of GluN1-specific T cells on day 42, which was abrogated by the anti-CD20, suggesting a contribution of GluN1-specific CD20+ T cells in B cell activation (Supplementary Fig. 4C).

### Brain-bound IgG from NMDAR mice precipitates NMDAR

Quantitation of the IgG bound to brain was performed by immunohistochemistry on tissue obtained at days 42 and 71 in the indicated subsets of NMDAR mice and controls. Compared to controls, NMDAR mice had a significant increase of brain IgG at days 42 and 71, which was substantially decreased or not significant in mice treated with anti-CD20, but not NMDAR-PAM (Supplementary Fig. 5A-C). The absent effect of NMDAR-PAM on IgG binding was in line with previous reports that used a mouse model of passive transfer of patients’ NMDAR antibodies, in which NMDAR-PAM antagonized and reversed the pathogenic effect of antibodies without affecting antibody binding to NMDAR.^18, 19^

The specificity of the brain-bound IgG for NMDAR was confirmed by precipitation of brain IgG, which co-precipitated NMDAR (Supplementary Fig. 5D).

### NMDAR mice show a decrease of NMDARs and impaired hippocampal plasticity, reversible with treatment

Having shown that NMDAR mice develop an anti-NMDAR-specific immune response and that the associated antibodies bind to brain NMDARs, we determined whether the neuronal surface content of NMDAR clusters was modified in the cerebral cortex and hippocampus of mice examined at days 42 and 71 (Fig. 2A). Confocal quantification of NMDAR clusters showed a significant reduction of total and synaptic surface clusters at day 71 in the hippocampus, dentate, and cerebral cortex of NMDAR mice, but not in controls (Fig. 2B-G). In the hippocampus, the reduction of total and synaptic NMDAR clusters was detectable at day 42 (Fig. 2B-C).

**Figure 2:**
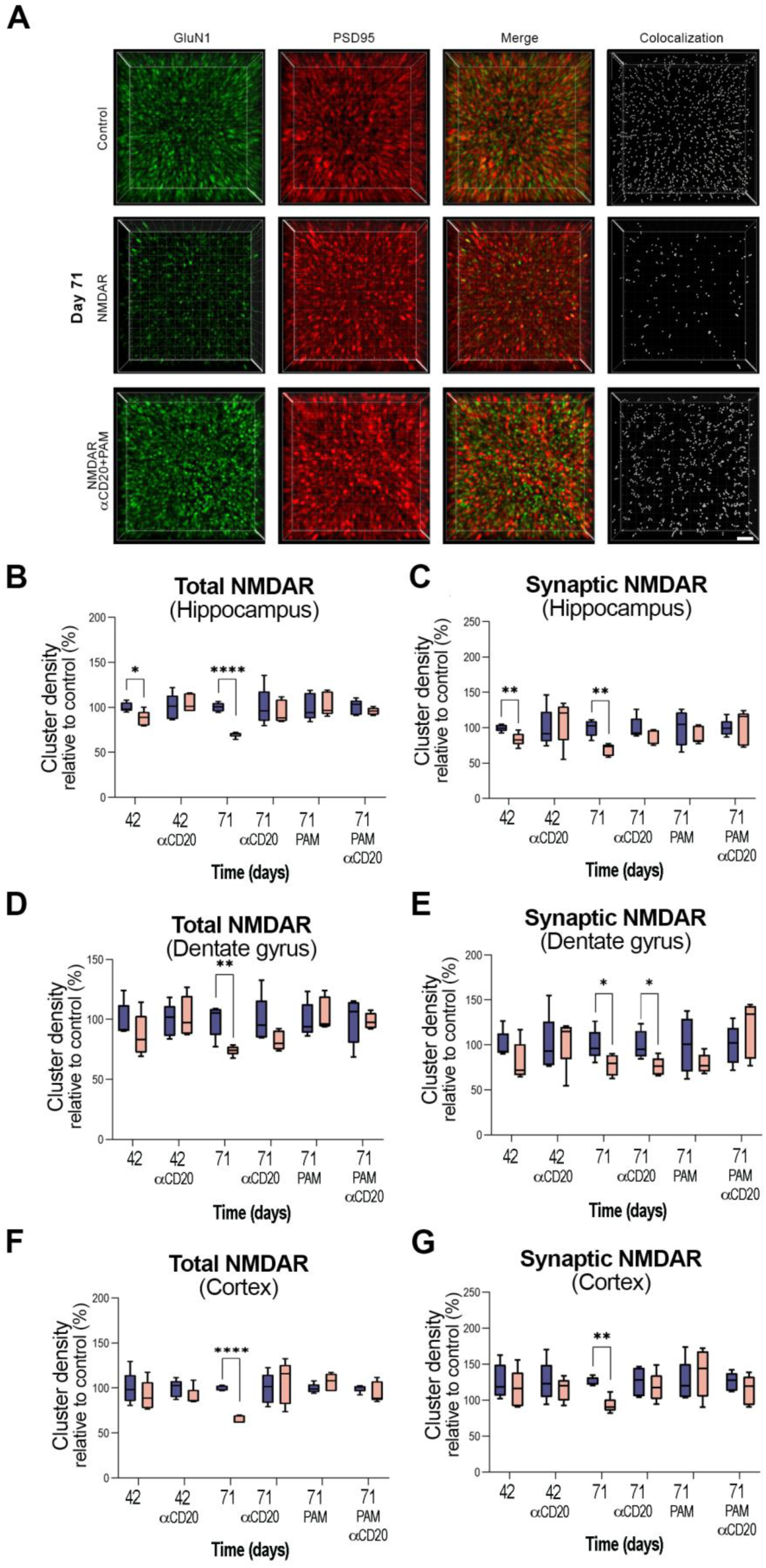
Treatments with anti-CD20 and NMDAR-PAM restore the levels of NMDARs in NMDAR mice. **(A)** 3D projection and analysis of the density of total cell-surface NMDAR (GluN1) clusters, PSD95, and synaptic NMDAR clusters (defined as those that co-localized with PSD95) in a representative area of CA1 of the hippocampus at day 71. Top row corresponds to the CA1 region of a control mouse; middle row, NMDAR mouse; lower row, NMDAR mouse treated with anti-CD20 and a positive allosteric modulator (PAM, SGE301) of NMDAR. Merged images were postprocessed and used to calculate the density of clusters (density = spots/µm^3^). Scale bar = 2 µm. For each animal 42 square images similar to those shown in A were examined (9 from CA1, 9 from CA3, 9 from dentate, and 15 from cortex). Quantification of the density of NMDAR clusters showing **(B)** total and **(C)** synaptic NMDAR clusters in a pooled analysis of hippocampal areas (CA1, CA3), **(D)** total and **(E)** synaptic NMDAR clusters in the dentate gyrus, and **(F)** total and **(G)** synaptic NMDAR clusters in the cortex. For the 3 brain regions assessed, untreated NMDAR mice show a significant reduction of synaptic and extrasynaptic clusters on day 71, and substantial or significant reduction of clusters on day 42. Treatment with anti-CD20, NMDAR-PAM, or both combined, restored the levels of NMDAR, but on day 71 NMDAR mice only treated with anti-CD20 started showing a new reduction of NMDAR (significant in dentate gryus) coinciding with B cell repopulation (see Fig 4C). Mean density of clusters in control conditions was defined as 100%. For each experimental condition 5 controls (blue) and 5 NMDAR mice (pink) were examined. Box plots show the median, and 25th and 75th percentile; whiskers indicate the minimum or maximum values. Significance was assessed by a mixed nested model and a value of p < 0.05 was considered significant.

By contrast, all but one subset of NMDAR mice treated with anti-CD20, NMDAR-PAM, or both, showed unaltered levels of total and synaptic NMDAR clusters at days 42 and 71 compared to those of controls. The only subset of treated NMDAR mice that on day 71 had reduced synaptic NMDAR clusters (noted in the dentate) was the group that received anti-CD20 (Fig. 2D-E).

Studies of long-term potentiation (LTP) in the hippocampus showed that untreated NMDAR mice had a significant impairment of plasticity at days 42 and 71 compared to that of controls (Fig. 3A-D). By contrast, all but one subset of NMDAR mice treated with anti-CD20, NMDAR-PAM, or both, showed unaltered LTP at days 42 and 71 compared to controls (Fig. 3B and 3E-H). Similar to the analysis of NMDAR clusters, only the subset of NMDAR mice treated with anti-CD20 showed LTP impairment at day 71 (Fig. 3B, F). Presynaptic release probability, as assessed by paired-pulse facilitation,^18^ was unaffected (data not shown).

**Figure 3:**
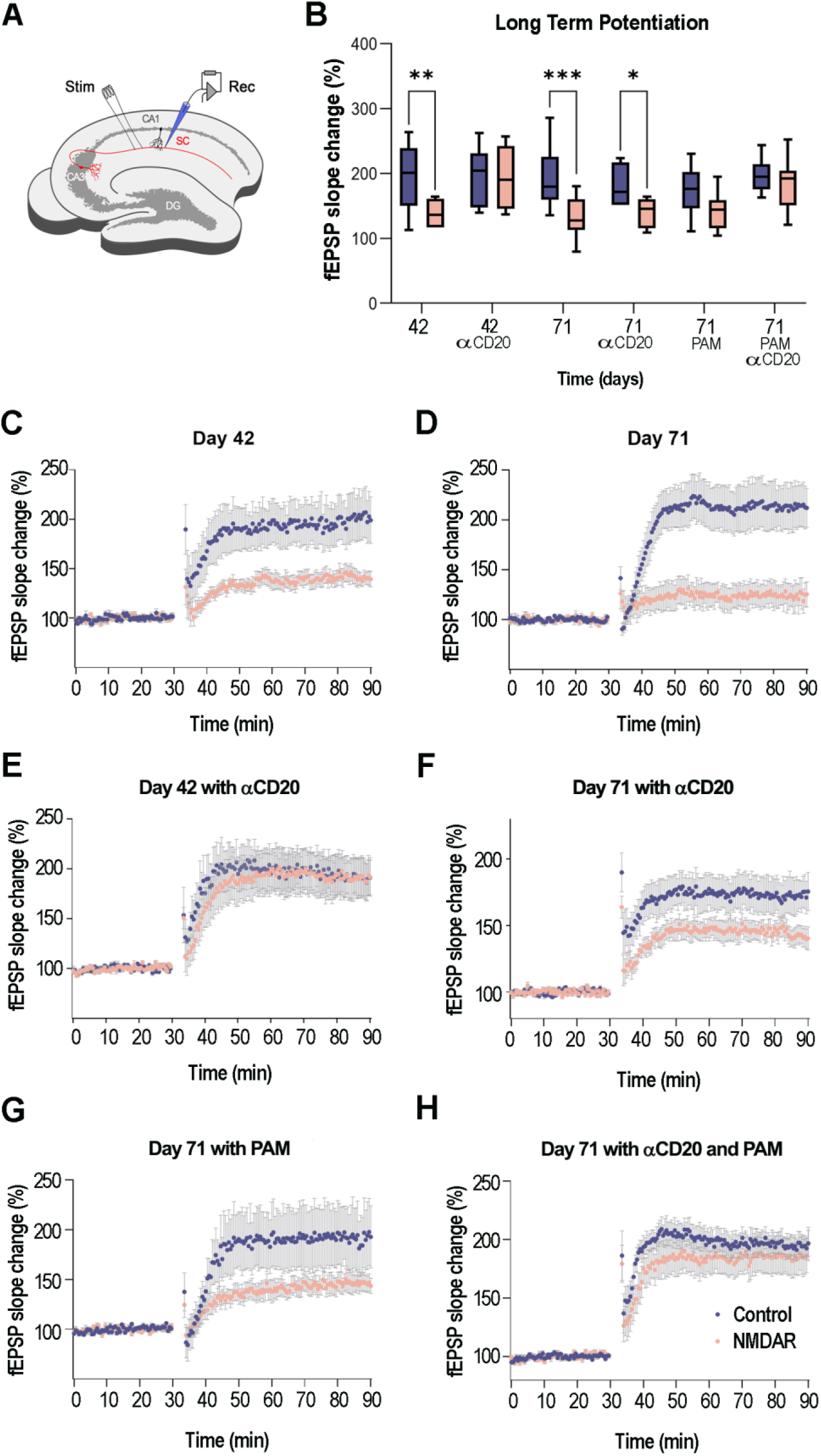
Hippocampal long-term potentiation (LTP) in NMDAR mice, and effects of different treatments. The Schaffer collateral pathway (SC, red) was stimulated (Stim) and field potentials were recorded (Rec) in the CA1 region of the hippocampus. LTP was induced by theta burst stimulation. **(B)** Compared with controls (blue), untreated NMDAR mice (pink) showed a significant reduction of field excitatory postsynaptic potential (fEPSP) slope change on days 42 and 71. NMDAR mice treated with anti-CD20 showed early recovery of fEPSP (day 42) but new reduction of fEPSP on day 71, coinciding with B-cell repopulation (see Fig. 4C). Box plot shows the median, 25^th^ and 75^th^ percentiles; whiskers indicate minimum and maximum values. Significance was assessed with a nested mixed model. A value of p < 0.05 was considered significant. **(C)** Time course of fEPSP recordings in control (blue) and NMDAR mice (pink) on day 42 (controls n = 6, NMDAR n = 7); **(D)** day 71 (controls n = 8, NMDAR n = 9); **(E)** day 42 treated with anti-CD20 (controls n = 6, NMDAR n = 6); **(F)** day 71 treated with anti-CD20 (controls n = 8, NMDAR n = 7); **(G)** day 71 treated with NMDAR-PAM (SGE-301) (controls n= 6, NMDAR n = 8); and **(H)** day 71 treated with both anti-CD20 and NMDAR-PAM (controls n = 6, NMDAR n = 6). The fEPSP values of all animals for each of the groups are presented as mean ± standard error of the mean (SEM).

These findings together with the indicated reduction of NMDAR clusters in cultured neurons exposed to IgG of NMDAR mice, suggest the NMDAR immune response causes a reduction of cell-surface NMDARs and that treatment with anti-CD20, NMDAR-PAM, or both, reverses this effect. Although anti-CD20 was effective in preventing all alterations at day 42, it was no longer as effective by day 71, coinciding with B cell repopulation (shown in the next section). The restoring of NMDAR cluster content in animals treated with NMDAR-PAM is similar to the reported effects of this PAM in cultured neurons exposed to patients’ NMDAR antibodies and in a model of passive transfer of patients’ antibodies.^18, 19^

### NMDAR mice have brain infiltrates of B and plasma cells with distinct treatment-response timings

The presence of brain immune cell infiltrates was determined by flow cytometry at days 42 and 71 in all subsets of NMDAR mice and controls (Fig. 4A). The brain of untreated NMDAR mice showed at days 42 and 71 a significant increase of pan-B cells (CD3-CD19+), memory B cells (CD3-CD19+CD27+), and plasma cells (CD19-CD138+) compared with the brain of untreated control mice (Fig. 4B-D). By contrast, the subsets of NMDAR mice treated with anti-CD20 showed at day 42 a significant reduction of brain pan-B cells and memory B cells, with cell counts not different from those of control mice, whereas by day 71 the presence of memory B cells had increased in NMDAR mice compared to controls (Fig. 4B-C). Conversely, the effects of anti-CD20 on brain plasma cells were not observed at day 42 (NMDAR mice had more brain plasma cells than controls), but by day 71 the number of plasma cells was markedly reduced in NMDAR mice and not different from that of controls (Fig. 4D).

**Figure 4:**
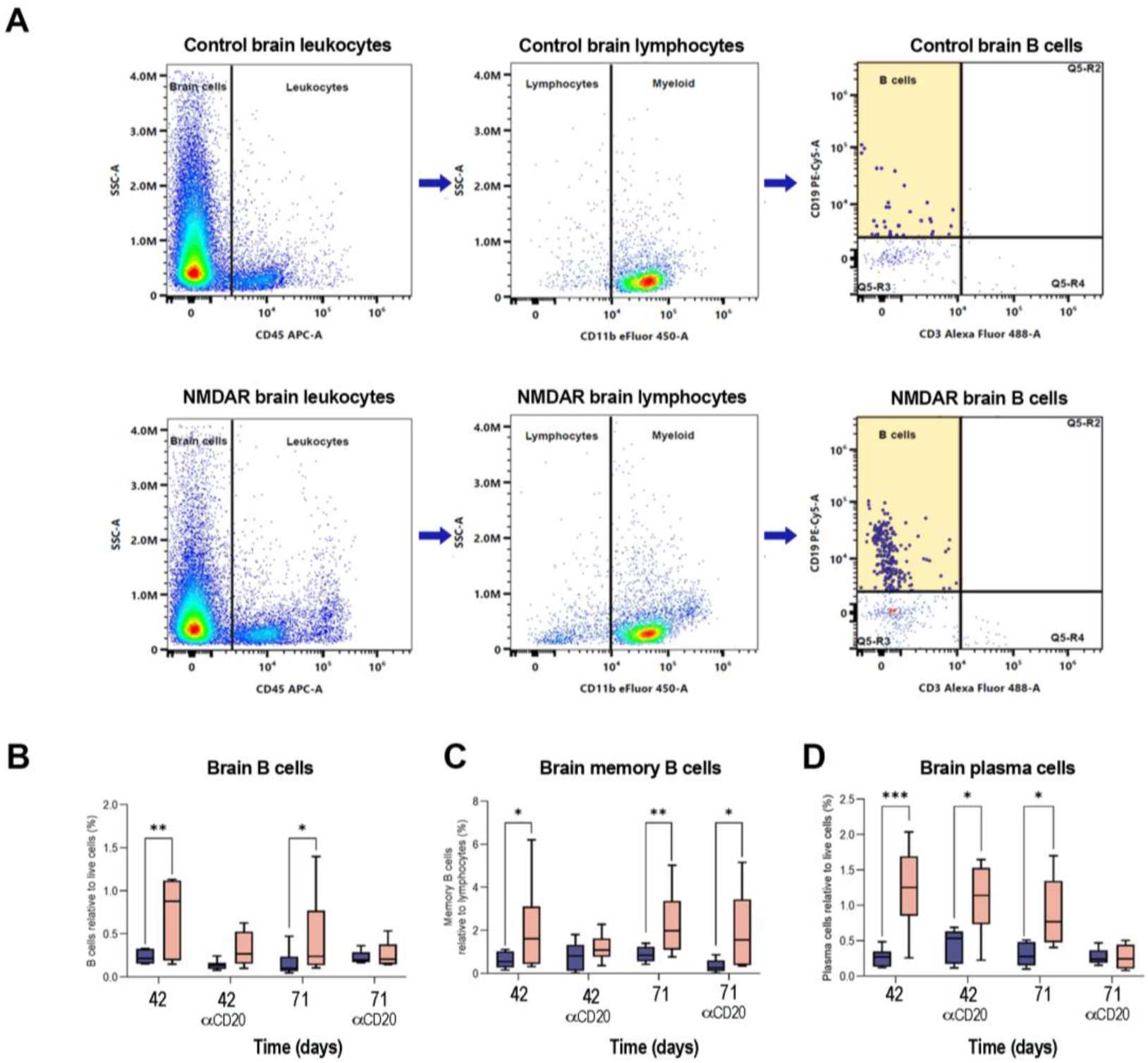
Presence of B cells and plasma cells in the brain of NMDAR mice, and effects of different treatments. **(A)** Flow cytometry scatter plots showing the presence of B cells and plasma cells in the brain of a representative NMDAR mouse and control mouse, illustrating the gating strategy to analyze the presence of the cell infiltrates. Live cells were gated for CD45 to select leukocytes and for CD11b to select lymphocytes. B cells were gated as CD3- and CD19+. Quantification of brain B cells **(B)**, memory B cells **(C),** and plasma cells **(D)** in control (blue, n=10) and NMDAR mice (pink, n=10) for all experimental conditions. Untreated NMDAR mice had a significant brain increase of total B cells, memory B cells, and plasma cells on days 42 and 71. NMDAR mice treated with anti-CD20 treatment showed a decrease of brain B cells on day 42 and plasma cells on day 71, but memory B cells started to repopulate on day 71. Box plot shows the median, 25^th^ and 75^th^ percentiles; whiskers indicate minimum and maximum values. Significance was assessed with a mixed model. A value of p < 0.05 was considered significant.

We did not find a substantial component of CD4 or CD8 T cells in the brain of NMDAR mice and controls by flow cytometry. Using brain immunohistochemistry, infrequent infiltrates of T cells (CD4>CD8) were identified in the meninges and brain parenchyma of both subsets of mice, with increased number of CD4+ T cells in NMDAR mice that was not significant (data not shown). Moreover, we did not find complement deposition in the brain of NMDAR mice (data not shown), which is similar to reported patients’ autopsy findings.^26, 27^

In parallel studies, antibody titres were determined in serum of mice before being sacrificed at days 42 and 71. In NMDAR mice, anti-CD20 alone or followed by PAM caused a partial reduction in serum NMDAR antibody titres on day 42, which was more marked on day 71. No reduction in antibody titres was observed in mice only treated with PAM (Supplementary Fig. 6). CSF from NMDAR mice treated with anti-CD20 showed less reactivity than CSF from untreated mice on day 42 and 71, although the reactivity was not completely abolished (Supplementary Fig. 6).

### NMDAR mice have brain microglial activation that co-localizes with IgG-bound to receptors

A significant co-localization of CD68 and Iba-1 (microglial activation) was noted on day 71 in the hippocampus of untreated NMDAR mice, and a similar trend was observed in the cerebral cortex (Fig. 5A-B). Microglial activation was not observed in subsets that received treatment with anti-CD20 or NMDAR-PAM. Because microglia might be involved in phagocytosis of NMDAR targeted by autoantibodies, we determined by confocal microscopy the triple co-localization of CD68, IgG and NMDAR in the hippocampus of untreated NMDAR mice and controls at day 71 (Fig. 5C). The findings showed a significant increase of this triple co-localization in the hippocampus of NMDAR mice (Fig. 5D-E). Super-resolution STED microscopy in several representative areas of triple co-localization demonstrated the presence of IgG and GluN1 in CD68-expressing endosomes of microglial cells, suggesting phagocytosis of antibody-targeted NMDARs (Fig. 5F-G). Overall, these findings support a pathogenic role of the microglia in the brain autoimmune process, likely contributing to the reduction of NMDARs.

**Figure 5:**
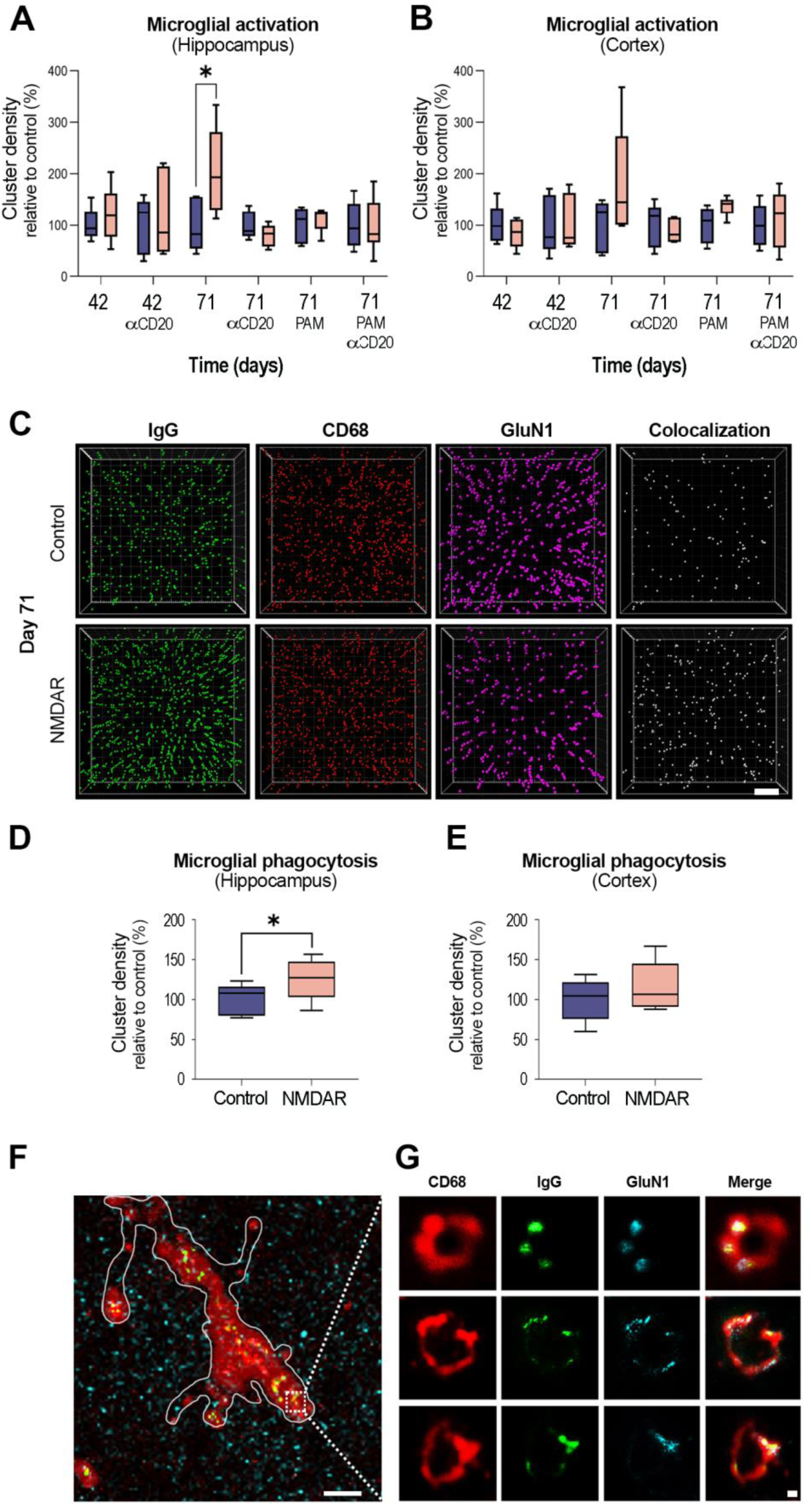
Microglia activation and phagocytosis of NMDARs bound to IgG. Quantification of microglia activation (defined as the co-localization of Iba-1 and CD68) in the hippocampus **(A)** and cortex **(B)** of NMDAR mice and controls; the areas of the hippocampus (CA1, Ca3, and dentate) and cortex are the same as those examined in Figure 2. **(C)** 3D projection and analysis of the density of clusters of IgG, CD68, total cell-surface GluN1, and their triple co-localization in a representative CA1 square region of an NMDAR and control mice at day 71. Merged images were postprocessed and used to calculate density clusters (density = spots/µm^3^). Scale bar = 2 µm. Quantification of triple co-localization of clusters (IgG, CD68, GluN1 = defined as microglial phagocytosis of NMDAR) in the indicated regions of hippocampus (**D**) and cortex **(E)** of untreated NMDAR and control mice on day 71. Mean density of clusters in control conditions was defined as 100%. For each experimental condition 5 controls (blue) and 5 NMDAR mice (pink) were examined. Box plots show the median, and 25th and 75th percentile; whiskers indicate the minimum or maximum values. Significance of microglia activation was assessed by a mixed nested model, and microglia phagocytosis by t-student. A value of p<0.05 was considered significant. Super-resolution imaging of a microglial cell **(F,G)** with Stimulated Emission Depletion (STED) microscopy shows that the triple co-localization (white) of CD68 (red), IgG (green), and GluN1 (cyan) occurs in the endosomes. **F** scale bar = 2 µm; **G** scale bar = 100 nm.

### NMDAR mice show multiple symptoms of anti-NMDAR encephalitis that respond to treatment

Compared to the corresponding controls, all four subsets of NMDAR mice (untreated, treated with anti-CD20, NMDAR-PAM, or both) developed a significant decrease of memory (Object Location index) that was detected at first evaluation (day 34) and persisted until the last evaluation (day 68) unless mice were treated (Fig. 6A). The subsets of mice that were treated with anti-CD20, NMDAR-PAM, or both, showed a significant improvement of memory at day 47 that was maintained until the last assessment (day 68) except for the subset that received anti-CD20 alone which showed relapsing memory impairment by the time of B cell repopulation (Fig. 6A). Because all mice subsets received from days 45-71 daily injections of NMDAR-PAM or vehicle, we also assessed the memory changes in NMDAR mice and controls that did not receive any treatment or injections. In these two groups the findings were similar to those in the groups of NMDAR mice and controls that received injections with vehicle alone, suggesting that the stress caused by daily injections did not affect memory (data not shown).

**Figure 6:**
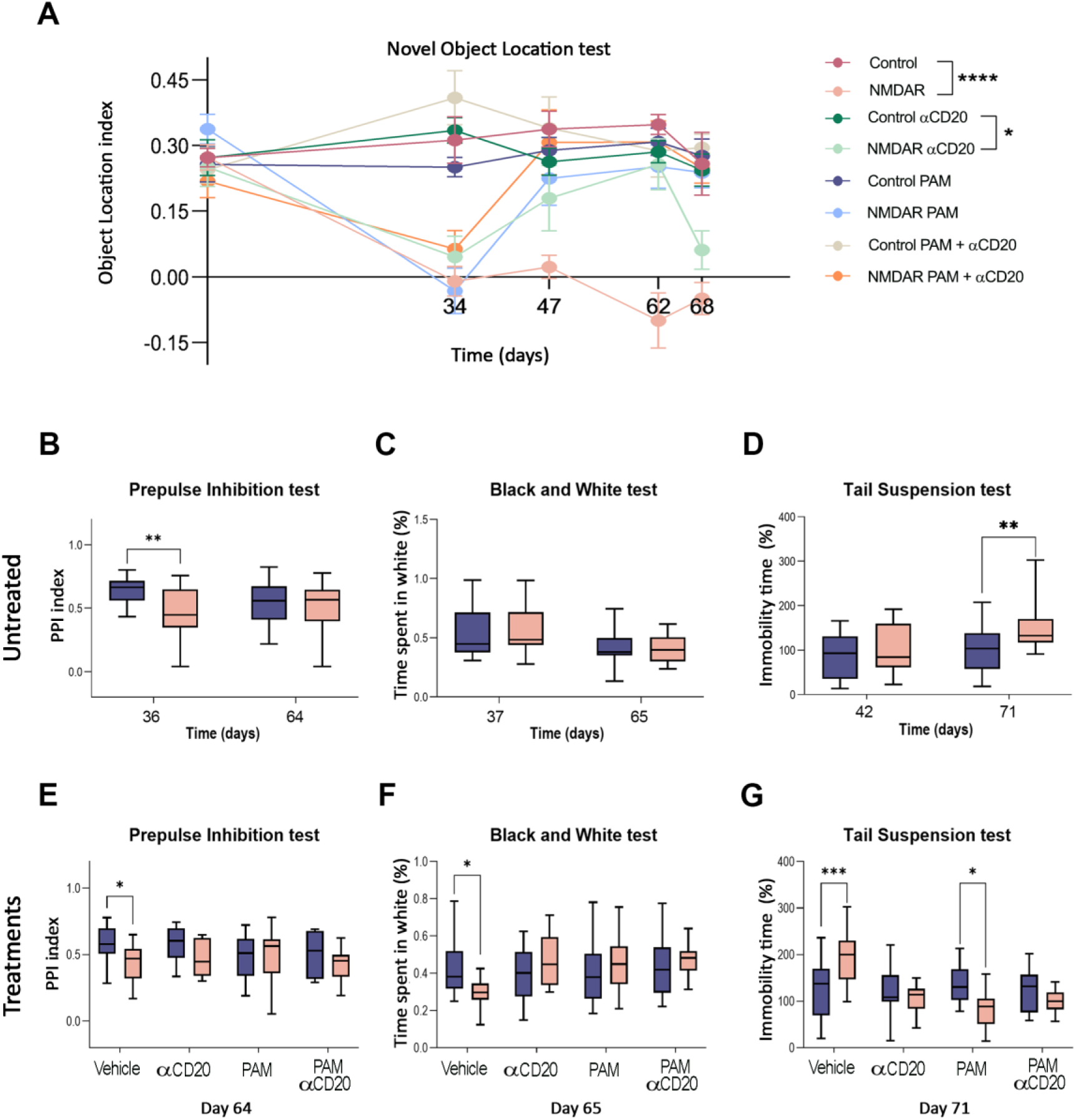
Behavioural and memory alterations in NMDAR mice, and effects of different treatments. (**A**) Memory assessment with the Novel Object Location (NOL) test in NMDAR mice and controls that received vehicle (no treatment), anti-CD20, NMDAR-PAM, and both treatments. On day 34, all NMDAR mice, but not controls, showed a significant decrease of memory. Treatment with anti-CD20 restored memory levels in NMDAR mice (days 47, 62), although by day 68, coinciding with B cell repopulation (see Fig. 4C), they showed relapsing memory impairment. NMDAR mice treated with NMDAR-PAM alone or the sequential administration of anti-CD20 and NMDAR-PAM showed improved memory from day 47 until the end of the study. Each experimental group included 12 animals. **(B-D)** Behavioural assessments in untreated NMDAR mice and controls that did not receive daily injections of vehicle: **(B)** compared to controls (blue), untreated NMDAR mice (pink) showed psychotic-like behaviour (prepulse inhibition test) on day 36, but not on day 64. Day 36, NMDAR mice n = 22, controls n = 23; day 64, NMDAR mice n = 23, controls n = 23. **(C)** Untreated NMDAR mice and controls showed similar levels of anxiety during the study (black and white test). Day 37, NMDAR mice n = 22, controls n = 23; day 65, NMDAR mice n = 24, controls n = 23. **(D)** Compared to controls, untreated NMDAR mice showed depressive-like behaviour (tail suspension test) on day 71, but not on day 42. Day 42, NMDAR mice n = 22, controls n = 23; day 71, NMDAR mice n = 24, controls n = 23. **(E-F)** Behavioural assessments (as in B-D) in NMDAR mice and controls that received no treatment (daily injection of vehicle) or one of the following treatments: anti-CD20, daily injection of NMDAR-PAM, or sequential combination of both treatments. Compared with controls (blue), untreated NMDAR mice (red) showed psychotic-like behaviour on day 64 **(E)**, anxiety on day 65 **(F)** and depressive-like behaviour on day 71 **(G)**. None of these alterations occurred in animals that received treatment with anti-CD20, NMDAR-PAM, or both. Each of the experimental groups in E-G included 12 animals. Box plots show the median, and the 25^th^ and 75^th^ percentile; whiskers indicate the minimum or maximum values. Line plots are represented as the mean ± SEM. Significance was assessed using a mixed model for the analysis of the tests of treated mice and two-way ANOVA with post hoc analysis, including multiple comparison corrections for the tests of untreated mice. A value of p < 0.05 was considered statistically significant.

In addition to memory impairment, untreated NMDAR mice developed early and transient psychotic-like behaviour (PPI test on day 36, but not day 64), and late depressive-like behaviour (TST on day 71, but not day 42) (Fig. 6B-D). Anxiety (BW test) was not affected in NMDAR mice or controls that did not receive any treatment or injections (Fig. 6C).

Since the administration of NMDAR-PAM was done with daily intraperitoneal injections that can result in stress and affect behavioural tests, the later assessment (as off day 45) of all paradigms of behaviour was controlled with subsets of mice that received daily injections of vehicle (Fig. 6E-G). Compared with these vehicle-injected controls, NMDAR mice showed psychotic-like behaviour (Fig. 6E), increased level of anxiety (Fig. 6F) and depressive like behaviour (Fig. 6G), which were all abrogated or improved by treatment with anti-CD20, NMDAR-PAM or both. These findings compared with the subset of untreated and not injected NMDAR mice which by the last assessment no longer have psychotic-like behaviour and did not show anxiety (Fig. 6B-D), suggest that NMDAR mice that received daily injections (either vehicle or treatment) had a potentiation or unmasking of NMDAR-related symptoms, which were successfully treated (psychotic-behaviour, anxiety) or improved (depressive-behaviour) with anti-CD20, NMDAR-PAM, and both.

In addition to memory and abnormal behaviours, 7 of 50 (14%) NMDAR mice, but not controls, exhibited motor stereotypies such as circling, self-biting, and walking backwards (Supplementary Video 2). None of these motor behaviours were observed after treatment with anti-CD20, NMDAR-PAM, or both. In addition, NMDAR mice showed a decrease in seizure threshold induced by pentylenetetrazol on day 42, which resulted in visible motor seizures (Supplementary Fig. 7). We did not find changes in locomotor activity in any subsets of NMDAR mice or controls, untreated or treated (data not shown). Altogether, NMDAR mice showed sequential psychiatric and neurologic alterations with persisting memory impairment, resembling the sequential clinical features of the human disease.

## Discussion

We introduce a mouse model of anti-NMDAR encephalitis that associates with a T cell dependent GluN1 IgG response and results in a reduction of cell-surface receptor content and NMDAR function, similar to the effects reported for antibodies of patients with anti-NMDAR encephalitis. To that end, we used a 30 amino acid peptide derived from the major GluN1 antigenic region of the human disease (containing amino acids N368/G369)^28^ in a novel immunization protocol that produced a robust synthesis of GluN1 polyclonal antibodies. These antibodies, predominantly of IgG1 subclass, targeted not only the immunizing GluN1 peptide but also epitope regions outside the peptide sequence, indicating epitope spreading.

These findings have not been previously investigated in models of anti-NMDAR encephalitis, and suggest a paradigm of immune-response different from that reported in myelin-oligodendrocyte glycoprotein (MOG)-experimental autoimmune encephalomyelitis (EAE) in which peptide immunization induces an EAE model that is mainly T cell mediated (B cell independent).^29, 30^ The longer GluN1 peptide length used in our model (30 amino acids), instead of shorter peptides (9-14 amino acid MOG fragments) used in the B cell independent EAE, together with the adjuvant AddaVax, which primes B cell responses may have played a role in the robust humoral response of our model. Indeed, AddaVax, a squalene-oil-in-water adjuvant, is known to be more effective in generating high antibody titres and CD4+T cell responses than Freund’s complete adjuvant or aluminium-based adjuvants^31–33^ and has fewer side-effects.^34^ Our studies with IFN-γ ELISpot on splenocytes from NMDAR mice confirmed the activation of GluN1-specific T cells, which was significantly decreased by the anti-CD20, suggesting an important contribution of GluN1-specific CD20+T cells in B cell activation, as reported in an EAE model.^35^

IgG isolated from NMDAR mice caused a reduction of NMDAR clusters and NMDAR-dependent calcium currents in cultured rat hippocampal neurons, similar to the alterations reported for the IgG of patients,^8, 24^ confirming that mice autoantibodies have direct effects on NMDARs. As a result, immunized mice, but not controls, showed NMDAR-specific brain-bound IgG, reduced content of synaptic and extrasynaptic NMDAR clusters, and significant impairment of hippocampal plasticity (LTP) similar to the alterations reported with passive cerebroventricular transfer of patients’ antibodies to mice.^9, 18^ However, different from the passive transfer model in which the duration of effects was shorter (∼14 days) until antibodies were cleared,^10, 18^ the current model showed NMDAR-related alterations for the entire duration of the study (71 days), providing the opportunity to assess different treatment strategies targeting at distinct disease mechanisms.

Another advantage of active immunization over passive transfer models is the possibility to study components of the immune response other than the antibody effects, such as brain inflammatory infiltrates, complement-mediated neuronal injury, and microglial activation. Analysis of brain inflammatory infiltrates, showed predominance of B cells and plasma cells, very infrequent T cells, absence of complement, and extensive microglial activation, overall resembling most of the findings reported in autopsies of patients.^1, 26, 27, 36^ A potential difference is that in patients, the frequency of T cells although low, might be higher than that observed in our model. Microglial activation is a consistent finding in patients’ autopsies,^1, 27^ suggesting it plays a pathogenic role.^37^ In the current study, NMDAR mice but not controls, showed a significant co-localization of CD68 (a phagocytic marker expressed by microglia/perivascular macrophages) with IgG and NMDARs. This triple co-localization, when assessed at the nanoscale level with super-resolution STED microscopy, was found to occur in endosomal/lysosomal structures, suggesting microglial phagocytosis of IgG-bound NMDARs.

Overall, these findings were accompanied by psychotic-like behaviour, memory deficits, depressive-like behaviour, variable presence of stereotyped movements (e.g., circling, self-biting, and walking backwards), and enhanced susceptibility to develop seizures (demonstrated with intracerebral electrodes). Interestingly, NMDAR mice not receiving treatment or injections of vehicle, developed psychotic-like behaviour earlier than depressive-like behaviour (as occurs in many patients),^5, 38^ whereas memory impairment persisted during the entire follow-up, and stereotyped movements occurred without stage preference. By contrast, psychotic-like behaviour remained detectable during the entire follow-up in untreated NMDAR mice stressed by daily (days 45-71) injections of vehicle.

The feasibility of the model to assess potential treatments, was tested with an anti-CD20 (equivalent to rituximab), and a synthetic analogue of 24(S)-hydroxycholesterol (SGE-301).^39^ SGE-301 is a potent and selective NMDAR-PAM that crosses the blood-brain-barrier and has been shown to antagonize and reverse the synaptic and behavioural alterations caused by patientś NMDAR antibodies in cultured neurons and passive transfer models.^18, 19, 25, 40^

NMDAR mice treated with the anti-CD20 showed rapid depletion of peripheral and brain B cell counts, accompanied by a decrease in brain-bound IgG, and recovery of NMDAR cluster density, hippocampal plasticity (LTP), and memory. These effects started wearing off about 5 weeks after treatment, when mice showed B cell repopulation and increased memory B cell infiltrates in the brain, accompanied by reduction of NMDAR clusters (initially detected in the DG), worsening synaptic plasticity, and return of memory impairment. These findings highlight the importance of B cell entry into the CNS, as suggested by neuropathological studies in patients,^27^ and a previous model examining the brain inflammatory infiltrates in untreated mice.^13^ Murine models of other disorders treated with anti-CD20 have shown variable duration of B cell depletion, ranging from 8 weeks post-treatment with three administrations of 200 µg of anti-CD20^41^ to 6 weeks after two administrations of 150 µg of anti-CD20.^42^ In our model, B cell repopulation occurred 5 weeks after a single 250 µg administration of anti-CD20, suggesting the depletion period is dose-dependent. These findings are in line with those in clinical practice which show the need of repeat cycles of rituximab to obtain therapeutic B cell depletion. The time lag between treatment administration and effects on symptoms and antibody levels is consistent with the findings in the EAE model in which symptom recovery associates better with B cell reduction than with the reduction of MOG antibody levels.^29^ Similarly, in our model, the decrease in brain plasma cells represented a delayed response, likely due to these cells not expressing CD20.

Of potential clinical interest, late daily treatment with NMDAR-PAM (SGE-301) restored NMDAR density, hippocampal LTP, and memory and behavioural functions, without modifying the levels of B cells or antibody synthesis. The mechanisms underlying these PAM effects are poorly understood, but previous studies by us and others showed that SGE-301 increased NMDAR function (e.g., open channel probability).^18, 39, 40^ In addition, single NMDAR molecule tracking in cultures of neurons exposed to patients’ NMDAR antibodies demonstrated that SGE-301 upregulated NMDAR surface diffusion in the post-synaptic compartment, which compensated for the antibody-mediated decrease in NMDAR surface dynamics and reduction of NMDAR clusters.^25^ Taken together, these findings suggest that SGE-301 or similar NMDAR-PAMs (e.g., some designed for oral bioavailability) could be an effective adjuvant treatment for anti-NMDAR encephalitis, particularly during the prolonged post-acute stage, when cognitive and psychiatric symptoms persist, and maintenance or escalation of immunotherapy may not be needed.^6^ In our model the early use of an anti-CD20 combined with a later administration of SGE-301 resulted in abrogation of all clinical and neurobiological alterations.

This study has several considerations and limitations. Because the model examines multiple clinical and biological paradigms, requiring multiple subsets of NMDAR mice and controls without and with several treatments, we designed the follow-up for 10 weeks after initial immunization. We chose this experimental design with the rationale that if active immunization recapitulated the clinical disease course, the model would provide important immune and neurobiological insights and be well positioned to test the effect of clinical standard care (anti-CD20) and an experimental NMDAR-PAM (SGE-301) that had been efficacious in a passive transfer model of patients’ antibodies. Our data strongly support this preclinical model of anti-NMDAR encephalitis as means to examine the course of the disease, and we believe the model can be followed for increasing amounts of time to investigate more long-term aspects of the disease. Indeed, further studies (not included here) in which we followed antibody titers for 4 months, showed that 8 of 10 untreated NMDAR mice remained with high antibody titers (similar to those of day 71); the other two animals only showed a mild spontaneous reduction of titers. Although we confirmed the presence of antibodies in CSF, the small amounts of CSF were a logistic problem that precluded several studies; thus, most investigations were performed with IgG isolated from serum. Mice showed propensity to develop seizures and status epilepticus, but not spontaneous seizures, which is a frequent feature in patients.^5^ The subsets of untreated mice with brain implanted electrodes were examined on day 42; we did not explore whether this lower seizure threshold could be treated with anti-CD20 or NMDAR-PAM, which is a task for the future. Finally, although changes in immunological, neurobiological, and behavioural paradigms usually occurred in parallel (e.g., B cell repopulation and re-emerging of neurobiological and memory problems), the correlation with serum antibody titers was not perfect, an observation also made in patients.^20^

Our current demonstration that GluN1 peptide immunization leads to a polyclonal NMDAR antibody response, microglial/macrophage phagocytosis of IgG-NMDAR complexes, and synthesis of NMDAR antibodies in deep cervical lymph nodes, together with reports showing synthesis of autoantibodies in deep cervical lymph nodes of some anti-NMDAR patients,^43^ and microglial phagocytosis of autoantibody-NMDAR complexes in mixed neuron/microglia cultures,^37^ suggest an immunological paradigm. After immunological activation at regional lymph nodes close to the immunization site, NMDAR antibodies produced systemically and by brain infiltrating B cells/plasma cells cause a reduction of neuronal NMDAR (as occur in the human disease)^8, 27^ accompanied by phagocytosis of NMDAR-IgG complexes by microglia/macrophages. These brain antigen-presenting cells likely contribute to epitope spreading by presenting new fragments of phagocytised NMDAR to CD4 T cells, resulting in a NMDAR-specific polyclonal B cell/ antibody response, probably at deep cervical lymph nodes.^44, 45^ Evidence of naïve T-cell priming and epitope spreading in the brain have been shown in models of EAE and other autoimmune encephalomyelitis.^46^

Our model can now be adapted in many ways, for example extending the follow-up, increasing the number of administrations of anti-CD20, using simultaneously anti-CD20 and NMDAR-PAM, or considering new therapies (e.g., CAR T cell technology). It also offers the opportunity to explore at the cellular and circuitry levels the alterations underlying the prolonged memory and behavioral changes, typical of the post-acute stage of anti-NMDAR encephalitis.^6, 47^ Finally, an important task for the future is to determine how a systemically triggered neuronal immune response (as occur in patients with teratoma) reaches the CNS, and the role of brain antigen-presenting cells (microglia/macrophages) and deep cervical lymph nodes in fine tuning the immune response (e.g., epitope spreading, antibody affinity).

## Acknowledgements

Moreover, we extend our gratitude to Mercedes Alba and Eva Caballero, Balma Serrano and Rafael Marín (FCRB-IDIBAPS), Maria Marsal, Merche Ribas, and Gustavo Castro (ICFO) for their technical support.

