## Supplementary material for "Neuro-immunobiology and treatment assessment in a mouse model of anti-NMDAR encephalitis"

### Supplementary methods

### Supplementary methods

#### Animal experiments

A total of 275 female C57BL6/J mice (Charles River) were used for all experiments. Sample size was calculated based on previously published studies using passive transfer of patients' antibodies and similar experiments.[1-3] Each mouse was considered as a experimental unit. Mice were randomly allocated to an experimental group based on a random number generator. Confounders were minimised by having two control and two NMDAR mice housed in the same cage for all experimental conditions. Investigators were blinded to the experimental allocation during behavioral and tissue investigations until data analysis.

#### Polyclonal antibody response and epitope spreading in NMDAR mice

Peptide and neuronal lysate immunoblot

The immunizing peptide $GluN1_{356-385}$ and hippocampal neuronal culture lysates were run in a gel, transferred to a nitrocellulose membrane (1704158, Bio-Rad, Hercules, CA), and incubated with pooled serum from 5 representative NMDAR mice (1:200) intact or pre-absorbed with the peptide (0.25 µg/µl) for 2 h at room temperature (RT). The reactivity was developed following a standard enhanced chemiluminescence developing kit (RPN2108, GE Healthcare, Chicago, IL).

Cell-based assays using GluN1 mutants

The GluN1 mutants have been previously described[4, 5] and are shown in Supplementary Fig. 1. In brief, G369I and G369S are single point mutations where glycine 369 was replaced by an isoleucine or serine, respectively. The "top lobe construct" carries a deletion of residues 26-140 and 275-349 in the top lobe of the amino-terminal domain (ATD). The deleted-construct carries a deletion of residues 12-385, containing all amino acids present in the immunizing peptide GluN1$_{356-385}$. HEK cells transfected with these mutants were used for immunocytochemistry following the same cell-based assay and serum dilutions used to test mouse samples with native GluN1. Briefly, HEK293 cells transfected with GluN1/GluN2b in equimolar ratios, or the indicated mutants, were grown for 24h after transfection. All cells were routinely grown in the presence of ketamine (500 µM) to prevent cell death after transfection. Transfected cells were then fixed with 4% paraformaldehyde for 5 min at RT, permeabilized with 0.3% Triton X-100 for 5 min at RT, and incubated with mouse serum diluted 1:40 and a commercial rabbit polyclonal antibody against the c-terminal region of GluN1 (dilution 1:5000, G8913, Sigma) overnight at 4 ºC. Cell were then washed with PBS, and incubated with the corresponding fluorescence secondary antibodies (Alexa Fluor 488 goat anti-mouse and Alexa Fluor 594 goat anti-rabbit, both diluted 1:1000 and from Invitrogen) for 1 h at RT.

**Determination of IgG subtype**

IgG subtype of antibodies from NMDAR mice was determined by CBA with HEK293 cells transfected with GluN1/GluN2b. Transfected cells were fixed and permeabilized as indicated above, and incubated with mouse serum diluted 1:40 and a commercial rabbit polyclonal antibody against GluN1 (dilution 1:5000, G8913, Sigma) overnight at 4 ºC. Cells were then washed with PBS and incubated with specific Alexa Fluor 488 goat anti-mouse IgG1 (A21121), IgG2a (A21131), IgG2b (A21141), or IgG3 (A21151) and Alexa Fluor 594 goat anti-rabbit (all diluted 1:1000 and from Invitrogen) for 1 h at RT.

**Pathogenicity of antibodies from NMDAR mice**

Cultures of primary hippocampal neurons

Primary hippocampal neurons were obtained from day 18 embryos of Wistar rats, as reported.[6] Dissociated neurons were seeded on coverslips and grown in Corning® 35 mm x 10 mm dishes (Sigma-Aldrich, St Louis, MI, US) containing 1 ml of Neurobasal medium + B-27 Supplement (ThermoFisher, Waltham, MA, US).

Calcium video microscopy

Calcium video microscopy was performed on 7-day *in vitro* primary cultures of dissociated rat hippocampal neurons transduced with the viral vector pAAV2-CAG-GCaMP5G at 2.5 · 1010 GC/mL as previously reported.[7] Five days after transduction, cells were treated with purified IgG from NMDAR mice or controls. One day later, cells were transferred to a chamber of an inverted fluorescent microscope equipped with a mercury lamp and a FITC filter cube (Eclipse TE20000-U, Nikon, Tokyo, Japan) and an ORCA flash4.0 v3 digital CMOS camera (Hamamatsu, Hamamatsu, Japan). The cell chamber was kept at 37°C with 5% CO2. Cells were treated with NBQX to block AMPA and KA receptors, and specifically visualize NMDAR dependent Ca2+ signal (fluorescence). A movie of 5 min with frames recorded every 100 ms was acquired. Shortly after starting the acquisition, 100 µM of NMDA and 1 µM of Glycine were added to the dish. The fluorescence signal over time was extracted by ImageJ.

Determination of effects of antibodies on NMDAR clusters in hippocampal neurons

To assess the effect of mice antibodies on NMDAR cluster density, serum from a pool of 5 NMDAR or control mice was added to the culture media (1:100) for 24 h. After removing the media and extensively washing with phosphate buffered saline (PBS), neurons were live incubated with human NMDAR IgG (1:200) for 30 min at room temperature to label the clusters on cell surface, as reported.[2] Subsequently, neurons were fixed with 4% PFA for 10 min, incubated with Alexa Fluor 488 goat anti-human IgG (1:1000, 109-545-088, Jackson ImmunoResearch, Newmarket, UK) for 1 h RT, and permeabilized with 0.3% Triton X-100 for 5 min. This was followed by incubation with a rabbit anti-PSD95 antibody (1:200, ab18258, Abcam, Cambridge, UK) for 1 h and subsequent incubation with Alexa Fluor 594 goat anti-rabbit IgG (1:1000, A-11012, Thermo Fisher Scientific). Cell surface clusters were captured using confocal microscopy (LSM710, Carl Zeiss, Jena, Germany). Images were deconvolved using Huygens Essential version 23.10 (Scientific Volume Imaging, The Netherlands) and quantified using Imaris 8.1 software (Oxford Instruments,).

**Determination of IgG deposits and precipitation of NMDAR-bound to IgG**

To determine the presence of mouse IgG in the brain, 5 μm-thick sagittal brain sections were blocked with 5% goat serum and immunostained for mouse IgG using Alexa Fluor 488 goat anti-mouse (1:500, A-11001, Thermo Fisher Scientific, Waltham, MA, USA) overnight at 4°C, as reported.[8] Slides were then mounted in ProLong Gold antifade (P36935, Thermo Fisher) and scanned under a Zeiss LSM710 confocal microscope (Carl Zeiss, Jena, Germany) with the EC-Plan NEOFLUAR CS 100x/1.3 NA oil objective.

To establish that the brain IgG was specifically bound to NMDARs, control and NMDAR mice brains were washed, homogenized in n-dodecyl-phoscholine 0.1% lysis buffer containing protease inhibitors (1:50, #P8340, Sigma-Aldrich) and ultracentrifuged (200,000 g). The supernatant was then incubated with protein A/G sepharose beads (20423, Thermo Fisher), precipitated, run in a gel, and blotted with a commercial GluN1 polyclonal rabbit antibody (G8913, 1:200, Sigma-Aldrich, St. Louis, MO, USA).

**Electrophysiology**

Electrophysiological studies on acute sections of mice brains were performed as reported.[2, 3] In brief, mice were deeply anesthetized with isoflurane and decapitated. Brains were removed in ice-cold, high sucrose extracellular artificial cerebrospinal fluid (aCSF1, in mM: 206 sucrose, 1.3 KCl, 1 CaCl2, 10 MgSO4, 26 NaHCO3, 11 glucose, 1.25 NaH2PO4; purged with 95% CO2/5% O2, pH 7.4), and subdivided into hemispheres. Thick (380 µm) coronal slices of hippocampus were obtained with a vibratome (VT1000S; Leica Microsystems, Wetzlar, Germany) and transferred into an incubation beaker with extracellular aCSF appropriate for neurophysiological recordings (aCSF2, in mM: 119 NaCl, 2.5 KCl, 2.5 CaCl2, 1.25 NaH2PO4, 1.5 MgSO4, 25 NaHCO3, 11 glucose, purged with 95% CO2/5% O2, pH 7.4). Slices were kept at 32°C for 1h and subsequently at RT for at least 1 additional h. For field potential measurements, single slices were then transferred into a measurement chamber perfused with aCSF2 at 2 ml/min at 28-30°C. A bipolar stimulation electrode (Platinum-Iridium stereotrode, PI2ST30.1A5, Science Products GmbH, Hofheim, Germany) was placed in the Schaffer collateral pathway. Recording electrodes were made with a puller (P-1000, Sutter Instrument Company, Novato, CA, USA) from thick-walled borosilicate glass with a diameter of 1.5 mm (BF150-86-10, Sutter Instrument). The recording electrode filled with aCSF2 was placed in the dendritic branching of the CA1 region for local field potential measurement (field excitatory postsynaptic potential, fEPSP). A stimulus isolation unit A385 (World Precision Instruments, Sarasota, FL, USA) was used to elicit stimulation currents between 25-700 µA. Before baseline recordings for long-term potentiation (LTP), input-output curves were recorded for each slice at 0.03 Hz. The stimulation current was then adjusted in each recording to evoke fEPSPs at which the slope was at 50-60% of maximally evoked fEPSP slope value. After baseline recording for 30 mins with 0.03 Hz, LTP was induced by theta-burst stimulation (TBS; 10 theta bursts of four pulses of 100 Hz with an interstimulus interval of 200 ms, repeated seven times with 0.03 Hz). After LTP induction, fEPSPs were recorded for 1 additional hour with 0.03 Hz. Recordings with unstable baseline measurements (variations higher than 20% in baseline fEPSPs) were discarded. Paired-pulse fEPSPs in the test pathway were measured before baseline recordings with an interstimulus interval of 50 ms. All recordings were amplified and stored using amplifier AxonClamp P2 (Molecular Devices, San José, CA, USA).  Traces were analyzed using Axon pClamp software (Molecular Devices, version 10.6).

**Determination of complement deposition in the brain**

To determine the presence of complement deposition, 5 µm-thick brain sections were permeabilized with 0.3% Triton X-100 for 10 min at RT and blocked with 5% goat serum. Then, slides were immunostained with a polyclonal rabbit anti-mouse C5b-9 antibody (1:200, ab55811, Abcam) for 2h at RT followed by the secondary antibody Alexa Fluor 488 goat anti-rabbit IgG for 1h at RT. Slides were then mounted in ProLong Gold antifade (P36935, Thermo Fisher) and scanned under a Zeiss LSM710 confocal microscope (Carl Zeiss, Jena, Germany) with the EC-Plan NEOFLUAR CS 100x/1.3 NA oil objective.

**Stimulated Emission Depletion (STED) microscopy**

To obtain a super-resolution image of microglia phagocytosis we utilized stimulated emission depletion (STED) microscopy. We conjugated the antibodies against GluN1 (human NMDAR IgG) with Alexa Fluor 594 (A20004, Thermo Fisher Scientific), and the antibodies against CD68 (MCA1957GA, Bio-Rad) with Abberior Star 635P (07679, Sigma-Aldrich). Alexa Fluor 488 goat anti-mouse (A-11001, Invitrogen, Carlsbad, CA, USA) was used to label mouse IgG.

Non-permeabilized brain sections were then sequentially incubated at 4ºC with the three indicated labelled antibodies (3 hours each at the following dilutions: 1:100 for Alexa Fluor

488 goat anti-mouse IgG; 1:10 for Alexa Fluor 594 human NMDAR IgG; and 1:20 for Abberior Star 635P anti-CD68). After incubation, the slides were washed and mounted with ProLong Gold (P36930; Molecular Probes, Eugene, OR) and scanned under a gated-STED microscope (TCS-SP8 STED 3X; Leica Microsystems, Wetzlar, Germany). For the fluorophores Alexa Fluor 488, Alexa Fluor 594 and Abberior Star 635P, we used excitation lines of 488, 594 and 635nm, and depletion lines (STED) of 592, 660 and 775nm, respectively. Fluorescence light was collected with HyD SMD molecule detectors on an HC PL APO CS2 100×/1.40 OIL objective (Leica Microsystems).

**Determination of brain T cells by immunohistochemistry**

To determine the presence of T cells by brain tissue immunohistochemistry, 5 µm-thick brain sections were permeabilized with Triton X-100 for 10 min at RT and blocked with 5% goat serum. Then, slides were immunostained with rabbit anti-mouse CD3 (1:200, ab16669, Abcam), rabbit anti-mouse CD4 (1:200, bs-0766R, Bioss), or rabbit anti-mouse CD8a (1:200, PA5-81344, Invitrogen) over night at 4ºC, and followed by the secondary antibody Alexa Fluor 488 goat anti-rabbit IgG for 1h at RT. Slides were then mounted in ProLong Gold antifade (P36935, Thermo Fisher) and scanned under a Zeiss LSM710 confocal microscope (Carl Zeiss, Jena, Germany) with the EC-Plan NEOFLUAR CS 100x/1.3 NA oil objective.

**Isolation of splenocytes**

Spleen from euthanized mice were harvested and placed in HBSS buffer (w/o Ca2+ and Mg2+, 14175-053, Thermo Fisher Scientific). The spleens were then minced and passed through a 70 µm cell strainer to create a single-cell suspension. The cell suspension was layered over Ficoll-Plaque PLUS (Cytiva, GE17-1440-02, Sigma Aldrich) and centrifuged at 800 g during 20 min without brakes. The mononuclear cell layer was collected and washed with HBSS buffer.

### Behavioural tasks

Novel Object Location (NOL) test

NOL test is a paradigm to study memory. Animals were habituated to an empty, squared arena (45x45 cm, Panlab, Spain) with visual cues, and underwent two daily trials of 15 mins each, for four days. The day of the test, animals were placed into the arena in presence of two equal objects positioned at two opposite corners and they were allowed to explore the objects for 9 min (familiarization phase). After a retention time of 3 h, animals were reintroduced to the arena, where one of the objects had been moved to a different corner. The animal was allowed to explore the objects for 9 mins (test phase) and the time of exploration of each object was recorded. A discrimination index (NOL Index) was calculated using the following formula: Time of exploration of the moved object minus time of exploration of the not moved object, divided by total time of exploration of both objects. A higher discrimination index indicates a better memory of the position of both objects. Object exploration is defined as any exploratory behaviour triggered by the presence of the object (sniffing, biting, touching) with the orientation of the nose toward the object within a distance of < 2 cm.[2]

Locomotor Activity (LA) test

Mice were assessed in LA boxes (9x20x11 cm, Imetronic, Passac, France), equipped with 2 rows of photocell detectors and placed in a low-luminosity environment (20-25 lux), as previously described.[9] Mice locomotor activity was recorded for 1 hour as horizontal activity and vertical activity.

Pre-pulse inhibition of the acoustic startle response (PPI) test

The PPI test is a classic paradigm to measure alternations in sensorimotor gating, which have been shown to occur in models of psychotic-like behavior.[10] Mice were habituated to being restrained using a plexiglass cylinder within a startle box (Panlab, Barcelona) in the presence of white noise (60 dB) and light for 15 min, for 3 days. The day of the test, after 5 min of habituation, a series of 10 trials of pulses (8 KHz. 115 dB, 40 miliseconds [ms]) was administered and the startle response (SR) was measured for 1000 ms (intertrial interval 29 seconds) in order to establish the basal response. The animal was subsequently exposed to a total of 40 trials randomly administered and equally divided into 4 different stimuli: pulse alone (8 KHz, 115 dB, 40 ms), prepulse alone (10 KHz, 80 dB, 20 ms), prepulse-pulse, and no stimulus (absence of pulse). Both habituation and test were always performed in presence of background white noise (60 dB) and light. The amount of inhibition of the SR due to the administration of the prepulse prior to the pulse was calculated as:

$SR\ inhibition$ (%) = 1 - ($average\ SR\ following\ prepulse\text{-}pulse\ stimulus$ / $average\ SR\ following\ pulse\ alone\ stimulus$) $x$ 100

Black and White (BW) test

BW test is a classic paradigm to measure anxiety-like behaviour in rodents; it measures the conflict between the innate exploratory behaviour and natural aversion to brightly illuminated areas.[11] The box includes a small black compartment and a big white compartment separated by a connecting gate (LE810, Panlab, Spain). The compartments are independently illuminated: the white one with a white led (500 lux) and the black one with a red led (10 lux). At the start of the session, mice were placed in the back compartment, head facing a corner. The latency of first entry into the white compartment and section reached in each entry, together with time spent, squares crossed, and number of entries into both compartments were recorded, tracked by Smart 3.0 software (Panlab, Spain) and used to evaluate anxiety.

Tail Suspension Test (TST)

TST is a classic task to measure depressive-like behaviour in rodents.[12] Mice are suspended by the tail in the automated tail-suspension box (76-1227, Harvard Apparatus, Massachusetts, USA). During a 6 min interval, energy, power, and total time of immobility are recorded. Long periods of immobility are characteristic of a depressive-like state.

**Seizure susceptibility study**

Pentylenetetrazol-induced seizures

Mice were handled for 2-3 days (corresponding to 10-15 days after the immune boost) before being placed in a square box (33x33 cm) with shallow bedding and <100 lux indirect light. After a 10 min baseline, the GABAaR antagonist pentylenetetrazole (PTZ, 2 mg/mL in saline) was injected at a dose of 40 mg/kg, intraperitoneally. Actimetry was tracked using Smart 3.0 software over the entire 1h session. A second frontal video camera was used for behavioral labeling off-line. All spontaneous actions were carefully observed at x 0.25-0.5 playback speed, annotated, and their association with PTZ treatment determined using the pointwise mutual information value. Labels distinctive of PTZ (laying on belly, crawling, pivoting, crunch posture with splayed hindlimb, head nodding, pacing, myoclonic jerks, clonus with loss of posture, forelimb sitting, freezing) appeared in a non-random order, and could be grouped into three main stages of seizure severity: hypoactivity, periodic jerks, and tonic-clonic. *In vivo* recordings and cFos induction (see below) confirmed this behavioral classification, as well as the occurrence of time-locked ictal activity during myoclonic jerks and tonic-clonic epochs. Jerks consisted of a combined downward motion of the head, jerk of the body and upward tail extension, all emerging in less than a second. A typical tonic-clonic episode lasted several seconds and followed a sequence of high-frequency jerks that made the mouse fall on its side, followed by righting, forelimb clonus and freezing. The incidence of seizure stages in control and immunized mice was performed blindly during the 50 min following PTZ injection. We tested the accuracy of detecting seizures behaviorally by comparing video and electrophysiological data in 7 implanted mice, and obtained >95 % coincidence for jerks and 100 % for tonic-clonic epochs.

cFos immunostaining

90 min after PTZ injection, mice were anesthetized and transcardially perfused with ice-cold formaldehyde as previously described.[13] Coronal cryostat sections (25 um-thick) were stained free-floating. After blocking in 2% BSA, 5% fetal bovine serum and 0.1 % Tween-20 in PBS, sections were incubated in rabbit cFos antibody (1:2000 dilution, ABE-457, Millipore) overnight at 4 ºC. For visualization, we used tyramide-mediated amplification following manual instructions (Invitrogen). Briefly, after rinses, sections were incubated with goat poly-HRP anti-rabbit antibody for 1h at RT, rinsed again, and incubated in reaction solution containing Tyr-AlexaFluor555 diluted 1:50 for 10 min in the dark. Sections were then counterstained with mouse anti-NeuN antibody (1:500, Millipore) overnight, and visualized with goat anti-mouse AlexaFluor488 (1:500, Invitrogen) for 2 h at RT. Epifluorescent images were acquired within linear range and exposure parameters fixed across groups and brain regions. Changes in cFos levels in neurons were analyzed with FIJI. NeuN staining was thresholded using the Moments function, a selection was then created and applied to the cFos channel to measure the grayscale intensity within. The cFos/NeuN signal ratio was normalized to that in vehicle-injected mice to generate a cFos matrix according to seizure severity using GraphPad 10.

Electrophysiological recordings in vivo

A bundle of 10-14 recording wires (Sandvik #PX000003) was glued onto a thin metal stick on one end, and cut to protrude 2.5-3 mm. On the other end, wires were stripped and individually wound to the pins of a 16-channel female connector (Omnetics #NSD-18-VV-GS). A silver wire (A-M systems #786000) was soldered to the ground pin. A drop of silver paint (RS Pro #123-9911) was applied to all wired pins, and then covered with abundant silicone glue (Dow #3145 RTV). The distal bundle tip was gold-plated to bring the wire impedance down to 100 KOhms. Briefly, the recording assembly was connected to a NanoZ kit (Neuralynx), tips were immersed in a diluted gold solution (final 2 % gold with 0.1% PEG; Neuralynx), and electroplated with repeated cycles of -0.05 uA to match progressively lower impedances.

In order to implant the recording drive in the right dorsal hippocampus, mice were anesthetized with isofluorane, immobilized in a stereotactic apparatus, and injected subcutaneously with bupivacaine (2 mg/kg). A 1 mm craniotomy was made, and the drive was slowly inserted (mm from bregma: -2 AP, 2 ML, -2 DV). The drive was secured with Superbond cement, and skin sutured. Mice received post-operative analgesia (meloxicam 5 mg/kg) for the next 3 days, and allowed to recover for a week before recording in a noise- and sound-insulated square box. Mice were tethered and allowed to freely move inside the box while signal was continuously acquired. Electrical signals were pre-amplified by connecting the drive to a headstage and acquired with an openephys board. A Bpod system connected to a PC was used to control and synchronize the videocamera (33 fps) and the electrophysiological signal with transistor-transistor logic (TTL) synchronization. Signals were analyzed in Python (3.7). They were first down sampled to 1 kHz, bandpass filtered between 0.1-300 Hz, and line noise removed using the notch filter.

### Supplementary figures

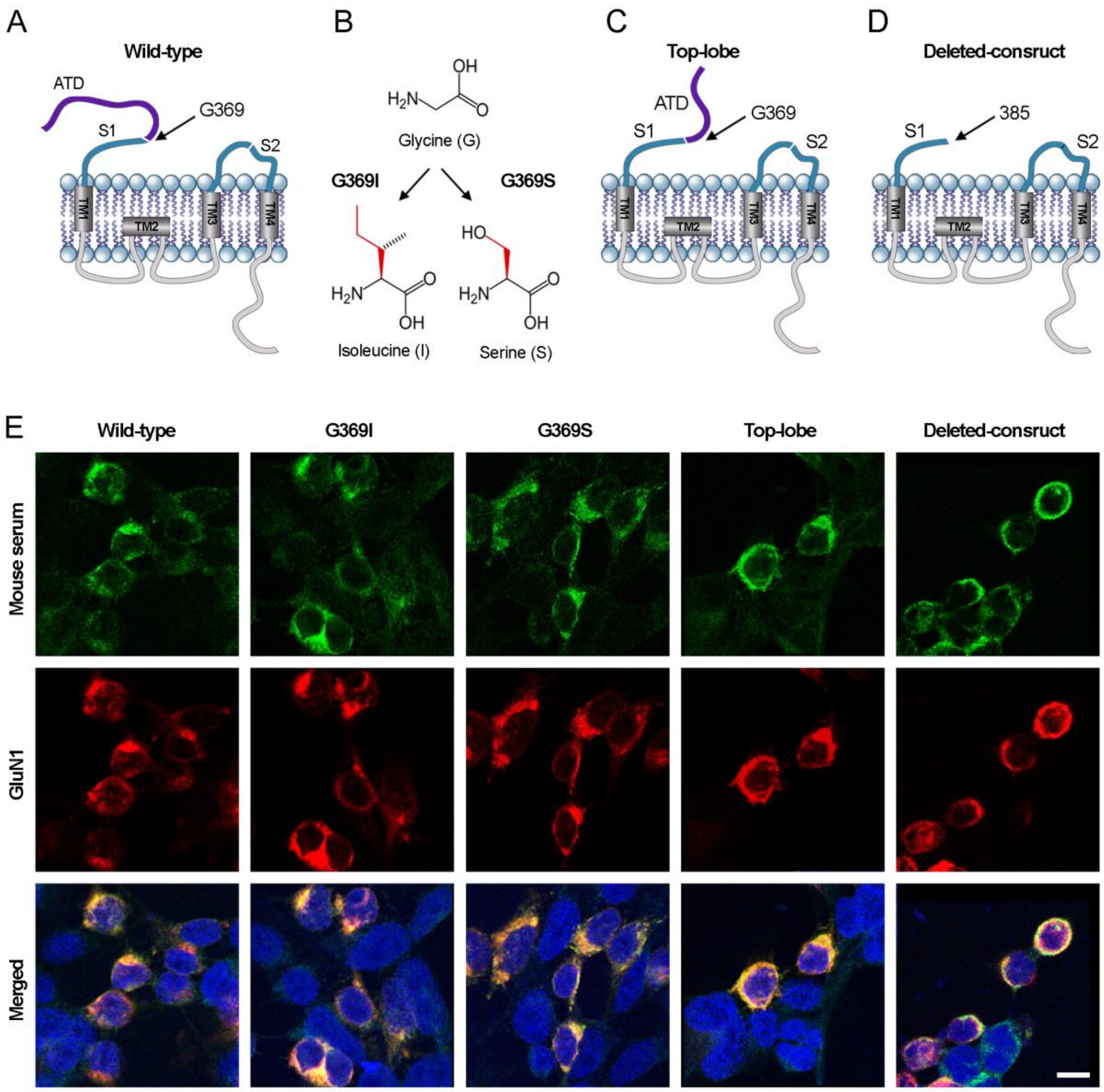

**Supplementary Figure 1: Mice immunized with GluN1$_{356\text{-}385}$ peptide produce polyclonal antibodies beyond the epitope region of the immunizing peptide**

Schematic representation of **(A)** native GluN1, **(B)** G369I and G369S, **(C)** amino terminal domain (ATD) top-lobe deleted, and **(D)** ATD deleted mutant constructs. **(E)** Cell-based assays with HEK293 cells expressing the indicated native or mutant GluN1 constructs and GluN2b, showing reactivity with serum of a representative NMDAR mouse (green), which colocalizes (yellow) with the reactivity of a commercial GluN1 antibody (red). All mice serum tested (n=10), but not controls, showed reactivity with all GluN1 constructs (not shown).

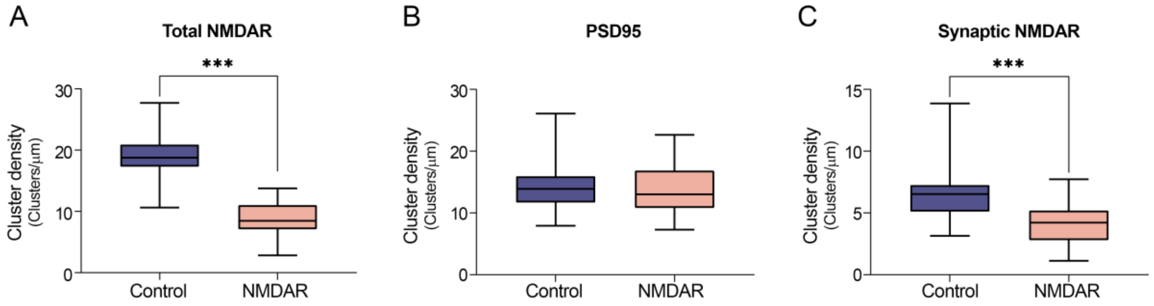

**Supplementary Figure 2: NMDAR mice serum cause a reduction of cell-surface NMDAR clusters in cultured rat hippocampal neurons**

**(A)** Total (synaptic and extrasynaptic) NMDAR clusters, **(B)** PSD95, and **(C)** synaptic NMDAR cluster density in cultures of rat dissociated hippocampal neurons treated with IgG purified from pooled serum of 5 control and 5 NMDAR mice. Results show a significant decrease in total and synaptic NMDAR cluster density in neurons treated with NMDAR IgG. 30 dendrites were analyzed per condition. Box plots show the median, 25th and 75th percentiles; whiskers indicate the minimum and maximum values. Significance was assessed using a t-test. A p-value of $< 0.05$ was considered statistically significant.

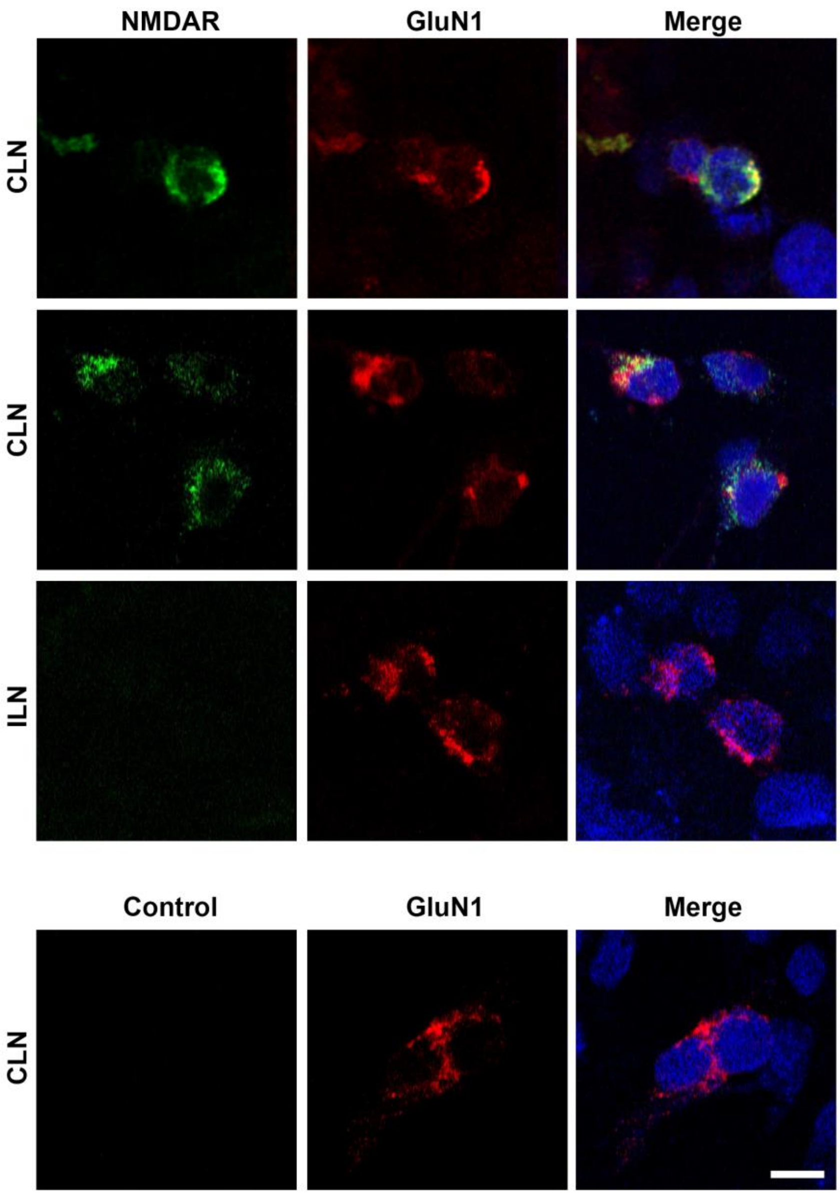

**Supplementary Figure 3: Cells from deep cervical lymph nodes from NMDAR mice produce GluN1 antibodies**

Cell-based assays with HEK293 cells expressing GluN1/GluN2b showing reactivity with antibodies secreted in the culture media by cells dissociated from deep cervical lymph nodes (CLN) of a representative NMDAR mouse (green). The reactivity of secreted antibodies colocalizes (yellow) with the reactivity of a commercial GluN1 antibody (red). Culture media of cells dissociated from inguinal lymph nodes (ILN) of the same NMDAR mouse, and CLN of a control mouse, do not show NMDAR reactivity.

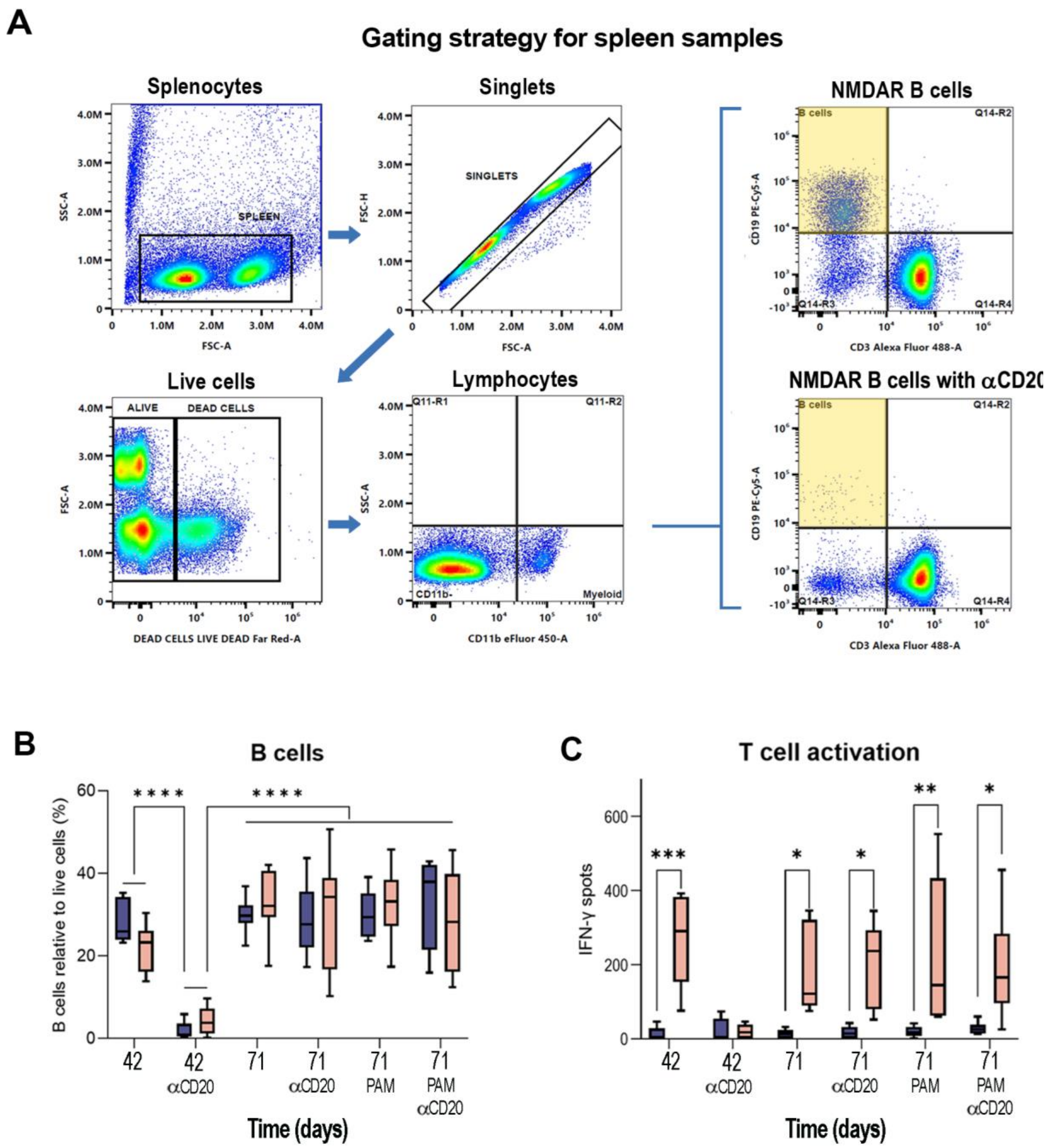
A
Gating strategy for spleen samples
Splenocytes
SPLEEN
SSC-A
FSC-A
Singlets
SINGLETS
FSC-H
FSC-A
Live cells
ALIVE
DEAD CELLS
FSC-A
DEAD CELLS LIVE DEAD Far Red-A
Lymphocytes
Q11-R1
Q11-R2
CD11b-
Myeloid
SSC-A
CD11b eFluor 450-A
NMDAR B cells
B cells
Q14-R2
Q14-R3
Q14-R4
CD19 PE-Cy5-A
CD3 Alexa Fluor 488-A
NMDAR B cells with αCD20
B cells
Q14-R2
Q14-R3
Q14-R4
CD19 PE-Cy5-A
CD3 Alexa Fluor 488-A
B
B cells
B cells relative to live cells (%)
Time (days)
42
42 αCD20
71
71 αCD20
71 PAM
71 PAM αCD20
C
T cell activation
IFN-γ spots
Time (days)
42
42 αCD20
71
71 αCD20
71 PAM
71 PAM αCD20

**Supplementary Figure 4: Effects of anti-CD20 on splenocytes and lymphocytes**

**(A)** Representative flow cytometry scatter plots illustrating the gating strategy used to identify B cells from spleen samples. Initial gaiting was performed on forward and side scatter properties to exclude debris. Subsequent gates were applied to select singlets and live cells. Next, cells negative for CD11b were selected as lymphocytes. B cells were gated as CD3- and CD19+ cells. Scatter plots show splenic B cells from a representative untreated NMDAR mouse and an NMDAR mouse treated with anti-CD20, exemplifying B cell depletion caused by anti-CD20. **(B)** Quantitation of splenic B cells in control (blue, n = 10) and NMDAR mice (pink, n = 10) under all experimental conditions. There is a significant decrease in B cells (day 42) in mice treated with anti-CD20, indicating effective B cell depletion, which by day 71 is no longer present (B cell repopulation) NMDAR-PAM (SGE-301) does not alter B cell number. **(C)** Quantitative analysis of the ELISpot experiment, indicative of IFN-γ production by splenocytes in response to GluN1$_{356–386}$ stimulation, in control (blue, n = 10) and NMDAR mice (pink, n = 10) under all experimental conditions. There is a significant increase of IFN-γ production in NMDAR mice, indicating T-cell activation, which is abrogated by anti-CD20 treatment on day 42. NMDAR-PAM does not alter T-cell activation. Box plots show the median, 25th and 75th percentiles; whiskers indicate the minimum and maximum values. Significance was assessed using a mixed model. A p-value of < 0.05 was considered statistically significant.

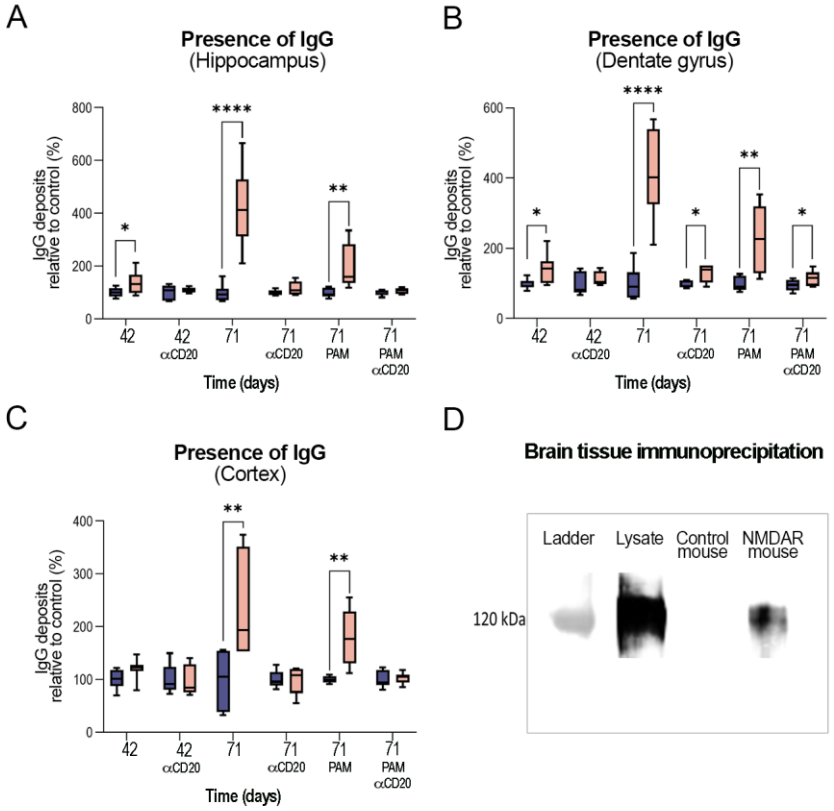

### Supplementary Figure 5: Presence of IgG bound to NMDAR in brain of NMDAR mice

Quantitation of mouse IgG brain deposits in controls (blue) and NMDAR mice (pink) across all experimental groups: **(A)** pooled analysis of hippocampal areas, **(B)** dentate gyrus, and **(C)** cortex. Compared to controls, NMDAR mice have a significant increase in IgG deposits, which is substantially decreased in mice treated with anti-CD20, but not NMDAR-PAM. For each experimental condition 5 controls and 5 NMDAR mice were examined. For each animal 42 square images similar to those indicated in Fig 2A were examined (9 from CA1, 9 from CA3, 9 from dentate, and 15 from cortex). Box plots show the median, 25th and 75th percentiles; whiskers indicate the minimum and maximum values. Significance was assessed using a nested mixed model. A value of $p < 0.05$ was considered statistically significant. **(D)** Immunoprecipitation of brain IgG in a representative NMDAR mouse and control, showing that in the NMDAR mouse the IgG is bound to NMDARs.

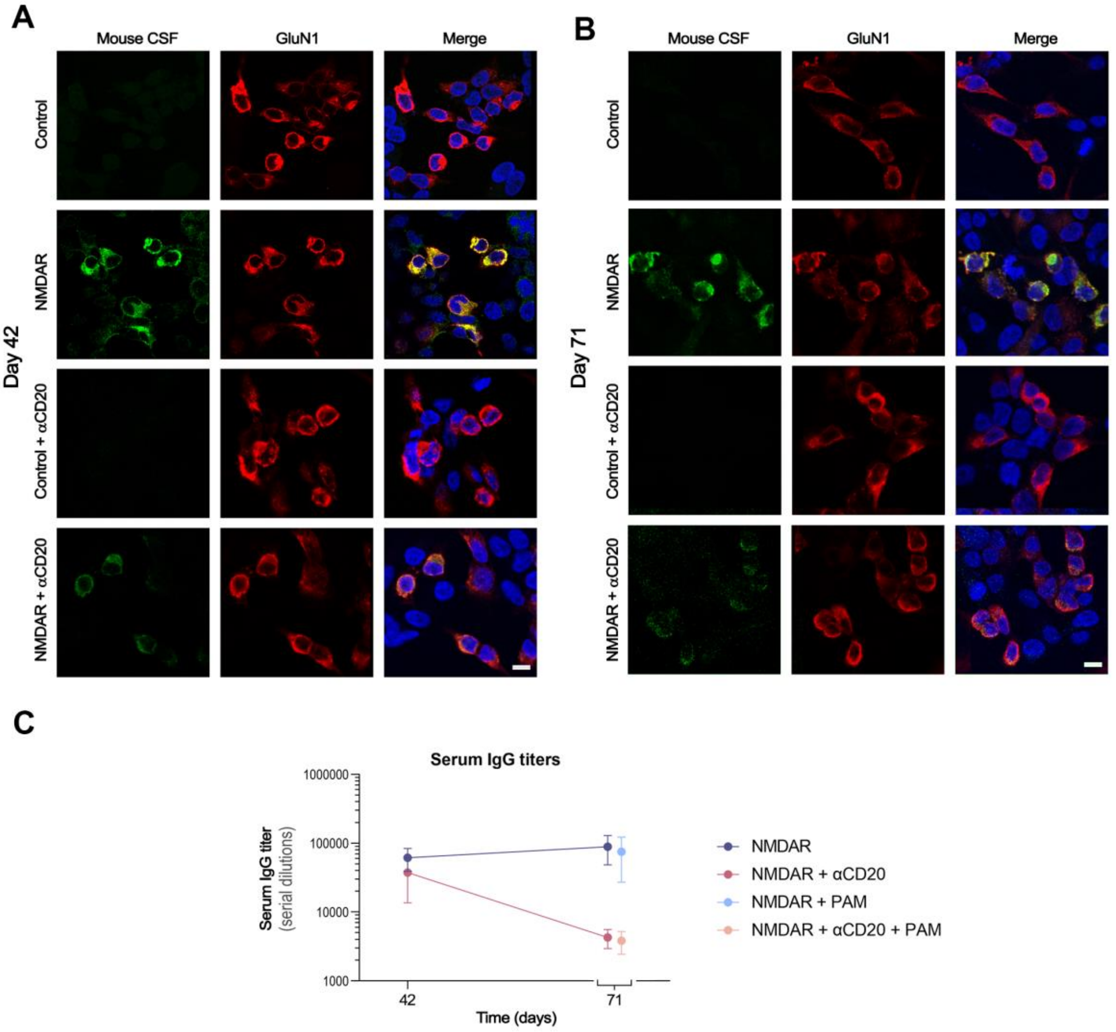

### Supplementary Figure 6: Effects of anti-CD20 on antibody titers

HEK293 cells expressing GluN1/GluN2b show reactivity with CSF from untreated NMDAR mice (green) at day 42 **(A)** and day 71 **(B)** after initial immunization. Mouse CSF reactivity colocalizes (yellow) with the reactivity of a commercial GluN1 antibody (red). CSF from NMDAR mice treated with anti-CD20 show a substantial decrease of immunoreactivity, although samples remain positive at both time-points (day 42 > day 71, the later barely visible). **(C)** Graphic representation of the change in serum titres (measured by CBA with serial serum dilutions) for all treatment groups of NMDAR mice (n = 6 for each experimental group). Anti-CD20 treatment reduces serum IgG titers by day 71. NMDAR PAM (SGE-301) does not alter IgG titers.

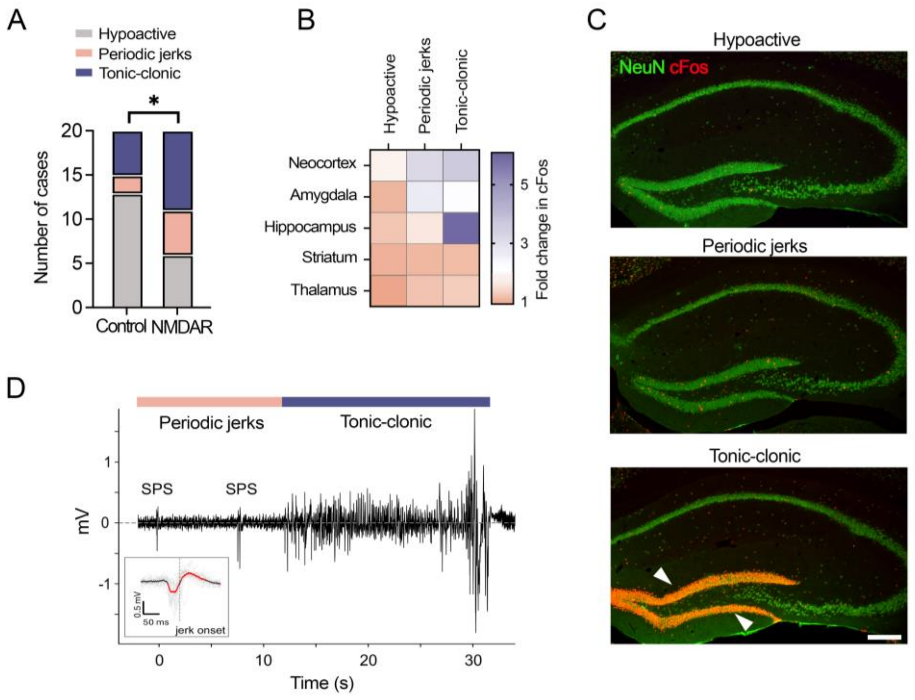

**Supplementary Figure 7: NMDAR mice show a decrease in seizure threshold induced by pentylenetetrazol**

(**A**) The distribution of maximal seizure stages shown in control and peptide immunized mice was quantified from data extending 1h after pentylenetetrazol (PTZ) injection (n=20/group). The fraction of mice showing generalized seizures (i.e. hypoactive versus jerks and tonic-clonic) was significantly larger in immunized mice, Chi square test, P = 0.026. (**B**) Quantification of cFos induction, a proxy of recent neural activation, across several brain regions in mice that reached different behavioral seizure stages (i.e. hypoactivity, myoclonic jerks, or continuous tonic-clonic seizures). Two-way ANOVA revealed an interaction between regional change in cFos and seizure stage, $F (8, 30) = 9.3$, $P<0.0001$ (n = 3 mice / group), indicating different patterns of neural activation depending on seizure stage. (**C**) Representative images of hippocampus double stained for cFos and the neural marker NeuN following different seizure stages. Arrows point to dentate gyrus granule cells where cFos induction was maximal after tonic-clonic seizures. Scale bar = 250 µm. (**D**) Example trace of the local field potential recorded in the hippocampus of a freely moving mouse. Sharp population spikes (SPS) were time locked with behavioral jerks, and transitioned to high-frequency spikes during the tonic-clonic episode. Inset, magnified average SPS aligned to jerk onset. Red points indicate voltage amplitudes significantly larger than baseline.

**Supplementary video**

**Supplementary Video 1: Reduction of NMDA-induced calcium currents in neurons treated with IgG from NMDAR mice**

Representative video of calcium imaging showing that neurons pre-treated with IgG from NMDAR mice show a significant reduction of NMDA-induced calcium influx compared to neurons pre-treated with IgG from controls. Scale bar = 10 µm.

**Supplementary Video 2: Motor stereotypies in NMDAR mice**

Video of four NMDAR mice showing different types of stereotypic movements and behaviours, such as walking backwards, self-biting, and circling.
